# Physiological mechanisms underlying coral acclimatization capacity to novel, multi-stressor conditions

**DOI:** 10.1101/2025.05.08.652802

**Authors:** Sarah L Solomon, Christian RA Lippens, Riccardo Mazza, Maya E Powell, Kelly W Johnson, Sophia Suvacarov, Rene M van der Zande, Verena Schoepf

**Affiliations:** Department of Freshwater and Marine Ecology, Institute for Biodiversity and Ecosystem Dynamics, University of Amsterdam, Amsterdam, the Netherlands; Environment, Ecology, and Energy Program, University of North Carolina at Chapel Hill, Chapel Hill, NC, 27514, USA

**Keywords:** Coral reefs, adaptive capacity, climate change, multiple stressors, reciprocal transplant experiment, extreme coral environments

## Abstract

Reef-building corals are under pressure to acclimatize and/or adapt to multiple co-occurring stressors, including climate change and declines in water quality. However, their capacity to do so is largely unknown. Yet, some corals naturally persist in habitats with multi-stressor conditions which can provide crucial insight into coral adaptive capacity under future ocean conditions. To assess the role of phenotypic plasticity and local adaptation in coral acclimatory responses to novel conditions, we reciprocally transplanted two coral species (*Siderastrea siderea*, branching *Porites*) between a multi-stressor, highly variable inland bay and a more stable fringing reef, and measured key phenotypic traits after 0, 4, and 12 months. Reef-origin corals demonstrated high plasticity when transplanted to the bay, as both species maintained high survival and calcification, and matched bay natives’ phenotypic traits. Yet, both species had reduced photosynthesis-to-respiration ratios, highlighting the risk of metabolic tradeoffs unless compensated for through increased heterotrophy. Conversely, bay-origin corals had enhanced photosynthetic performance but lower calcification on the reef. Reaction norms provided stronger evidence for environmental specialization in bay-origin than reef-origin corals, which could limit their use as stress-tolerant source corals for reef restoration. Overall, our findings suggest multi-stressor variability may promote specialized genotypes, rather than increased phenotypic plasticity.

## Introduction

As ecosystems face rapid climate change and environmental degradation globally, there is an urgent need to understand the adaptive capacity of organisms to increasingly extreme environmental conditions (Pacifici et al. 2015). Some organisms, such as tropical corals, are especially vulnerable as they are sensitive to multiple climate change stressors (i.e., ocean warming, acidification, and deoxygenation) (Hoegh-Guldberg et al. 2007; Altieri et al. 2017) as well as local stressors such as declines in coastal water quality (Nalley et al. 2021). Ocean warming and marine heatwaves disrupt the coral-algal symbiosis leading to coral bleaching and often large-scale mortality (Hoegh-Guldberg et al. 2023). In tandem, ocean acidification can reduce corals’ capacity to calcify and build the reef framework (Kornder et al. 2018; Cornwall et al. 2021). Ocean de-oxygenation and warming combined with the widespread degradation of coastal water quality has further led to an increase in localized low oxygen events, which are increasingly recognized as a threat to tropical coral reefs, particularly under climate change (Altieri et al. 2017; Pezner et al. 2023). As climate change intensifies and human population growth by coral reefs increases faster than global averages (Sing Wong et al. 2022), the future persistence of coral reefs depends on their capacity to rapidly adjust to multi-faceted environmental change. However, research to date has largely focused on coral capacity to acclimatize to one or two climate change stressors (Palumbi et al. 2014; Jury et al. 2024), and our knowledge on their adaptive capacity to multiple co-varying stressors and the physiological mechanisms facilitating acclimatization is limited (Camp et al. 2018; Guan et al. 2020; Schoepf et al. 2023).

Understanding how corals can acclimatize and/or adapt to multiple co-occurring stressors is a major challenge, as simulating these conditions *ex situ* over long time scales is difficult. However, some coral communities already persist naturally in environmentally extreme habitats, such as near CO_2_ vents, on macrotidal reefs or in semi-enclosed bays and mangrove lagoons and have provided crucial insights into the traits that facilitate success and survival at the edge of environmental limits (Camp et al. 2018; Burt et al. 2020; Schoepf et al. 2023). Notably, some of these extreme coral habitats are characterized by multiple co-varying stressors (Maggioni et al. 2021; de Jong et al. 2025), and can therefore serve as unique natural laboratories to investigate coral adaptive capacity to multiple stressors under ecologically realistic conditions (Camp 2022; Schoepf et al. 2023). Reciprocal transplant experiments, in particular, can offer insights into the mechanisms, time scales, and potential fitness trade-offs underlying adaptive capacity (Palumbi et al. 2014; Barott et al. 2021). To date, few studies have conducted such experiments in extreme coral environments, and most have focused on single stressors typically associated with climate change. For example, exposure of naïve populations to thermally fluctuating temperature regimes in back-reef pools, leading to significantly increased coral heat tolerance in some, but not all species, within <2 years (Palumbi et al. 2014; Klepac and Barshis 2020). Similarly, coral populations reciprocally transplanted for 17 months between naturally high- and low-pH habitats both exhibited fitness trade-offs under novel conditions, suggesting that local adaptation was underlying pH-tolerance (Barkley et al. 2017). While extreme coral environments with multiple co-varying stressors remain severely understudied, research from hot, acidified, low-oxygen mangrove lagoons has provided insight into potential fitness tradeoffs associated with living under multiple climate change stressors, with some studies reporting reduced calcification and higher metabolic demand (Camp et al. 2019; Jacquemont et al. 2022). However, coral adaptive capacity and potential associated fitness trade-offs under combined global and local stressors remain largely unknown.

Coral adaptive capacity to increasingly extreme conditions may be promoted by acclimatization via phenotypic plasticity utilizing their existing genetic repertoire, by genetic adaptation via evolutionary change across generations, or the interaction of both. Theory predicts that environmental heterogeneity should generally favor adaptive capacity via phenotypic plasticity, or “generalist” genotypes, while organisms from more stable environments may be locally adapted, or “specialists”, if there is enough selection for a given phenotype and gene flow across habitats is limited (Kawecki and Ebert 2004; Hereford 2009). Therefore, generalists should dominate in coral environments where multiple abiotic parameters naturally co-vary, as is the case in many coastal or back-reef habitats (Maggioni et al. 2021; de Jong et al. 2025). Reciprocal transplant experiments conducted across nearshore-offshore environmental gradients have demonstrated evidence of both generalist and specialist strategies in these populations (Kenkel et al. 2015; Baumann et al. 2021; Castillo et al. 2024). This highlights that coral responses to novel environmental conditions are highly context-specific, with more research needed to understand how the adaptive capacity of corals is influenced by intensity, heterogeneity, and different combinations of environmental stressors.

Here, we conducted a one-year reciprocal transplant experiment of two Caribbean coral species (massive *Siderastrea siderea* and branching *Porites* sp.) between a multi-stressor, environmentally variable inland bay and a more environmentally benign and stable fringing reef in Curaçao (southern Caribbean). Since the inland bay is characterized by warmer temperatures, higher turbidity and lower water quality, as well as strongly fluctuating temperatures, pH, and dissolved oxygen (DO) (Kuenen and Debrot 1995; de Jong et al. 2025), it is ideally suited to investigate coral acclimatization capacity to novel multi-stressor conditions that comprise both global and local stressors. We hypothesized that (1) corals from the bay habitat have high phenotypic plasticity and acclimatization capacity to the relatively more stable abiotic conditions on the fringing reef with no physiological fitness trade-offs – unless their plasticity is limited by environmental specialization or local adaptation, whereas (2) the acclimatization capacity of corals from the reef may be limited by local adaptation such that the novel, multi-stressor conditions in the bay may incur fitness costs. We measured key phenotypic traits after 0, 4, and 12 months of transplantation to assess plasticity, and further utilized proxies for fitness to estimate local adaptation of both bay and reef populations using reaction norms and the coefficient of selection against transplants (Kawecki and Ebert 2004; Hereford 2009).

## Methods

### Study sites, coral collection, and experimental design

The reciprocal transplant experiment was conducted between June 2022 and June 2023 on the leeward coast of Curaçao between a highly variable, multi-stressor inland bay, Spanish Water Bay (12°04’36.4” N, 68°51’35.8” W), and an environmentally more stable, clear-water fringing reef, Tugboat Reef (12°04’04.9” N, 68°51’42.7” W) (Fig. 1, S1). Spanish Water Bay is classified as an extreme *and* marginal coral environment (criteria defined in Schoepf et al. 2023) because (1) the mean and variance of multiple environmental variables deviate strongly from optimal conditions (Fig. 1E,F,G, Table S1) (Kuenen and Debrot 1995; de Jong et al. 2025) and (2) coral cover and species richness are much lower compared to nearby fringing reefs (Debrot et al. 1998; Vermeij et al. 2007; de Jong et al. 2025). The study sites have been previously described in detail (de Jong et al. 2025) and further details can be found in the Supplement.

**Fig. 1.**
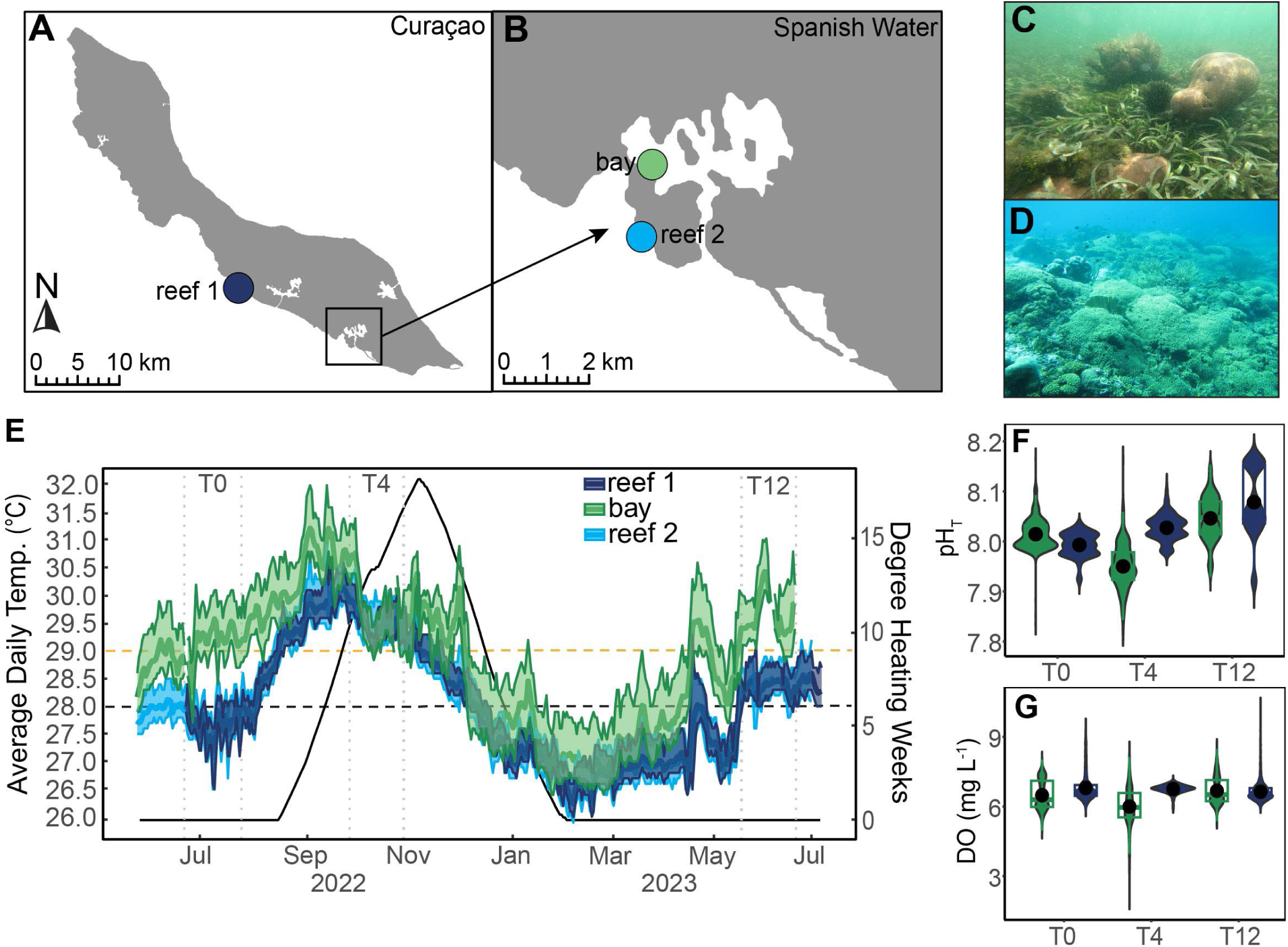
Map of Curaçao (A) and Spanish Water (B) showing bay (green) and reef (light blue) collection sites. Reef 1 (dark blue) in panel A is the location of the transplant reef, Carmabi House Reef. Photos highlighting typical benthic community structure at bay (C) and reef (D) collection sites. Temperature time series (average daily temperature, °C) and cumulative heat stress expressed as degree heating weeks (°C-weeks) across entire experiment (E; green = Spanish Water bay, dark blue = transplant reef 1/Carmabi House Reef, light blue = source reef 2/Tugboat Reef). Black dashed line = maximum monthly mean (MMM), yellow dashed line = bleaching threshold (MMM + 1°C). pH_T_ (F) and dissolved oxygen (DO) concentrations (mg L^-1^) (G) at the bay site and transplant reef following 0, 4, and 12 months of transplantation (T0, T4, T12)

Corals were collected between 5-9 June 2022. Parent colonies of massive *Siderastrea siderea* (n=10 genets per site) and branching *Porites* sp. (n=10 genets per site) were collected from each site using a hammer and chisel (n=40 total). Due to morphological plasticity characteristic of branching *Porites* in the Caribbean, we cannot be certain of the exact species (*P. divaricata*, *P. furcata*, or *P. porites*) but collected morphologies consistent with *P. furcata* (see Supplement for more details, Fig. S1). Bay-origin corals were collected from 1-2 m depth because few corals exist below 2.5 m due to higher turbidity. Reef-origin corals were collected from ∼3-8 m depth since the reef starts at ∼5 m, though most colonies were collected from ∼4-6 m depth. Despite differences in depth between bay and reef sites, light intensity differences were minimal due to the higher turbidity in the bay (Table S1).

After collection, each parent colony was fragmented into 5 fragments (ramets): one was immediately frozen at −80°C for baseline (T0) measurements, two were transplanted to their native habitat (to be collected after 4 and 12 months, respectively), and two were transplanted to their foreign habitat (to be collected after 4 and 12 months, respectively). Transplantation took place between 21–24 June 2022 (see Supplement for more details). This approach resulted in four transplant groups: 1) reef natives, 2) reef-to-bay transplants, 3) bay natives, and 4) bay-to-reef transplants. Corals were transplanted onto table-like structures at depths similar to their collection depths, corresponding to 5 m on the reef and 2.5 m in the bay. Corals collected from the reef (reef natives and reef-to-bay transplants) are referred to as “reef-origin corals”, while corals collected from the bay (bay natives and bay-to-reef transplants) are referred to as “bay-origin” corals.

Unexpectedly, corals from a separate pilot transplantation study experienced sudden partial mortality at Tugboat Reef in the days prior to the start of this experiment, possibly due to predation or a pathogen (see more info in the Supplement, Fig. S2). To avoid coral loss that would be unrelated to transplantation, we therefore changed the transplant site of the reef natives and bay-to-reef corals in this experiment to another fringing reef, Carmabi House Reef (12°07’14.4” N 68°58’11.8” W) (Fig. 1, Fig. S2). Comparison of environmental conditions at the two reef sites showed very similar averages and daily ranges for key parameters such as temperature, pH_T_, DO concentrations, photosynthetically active radiation (PAR), salinity, light and nutrients (Fig. 1E-G, Table S1), justifying the switch between reef collection and reef transplantation site.

### Environmental monitoring

Temperature was continuously monitored using data loggers between 28 May 2022 – 12 June 2023 at the bay and the two reef sites (10 min logging intervals). pH_T_, DO concentrations, PAR, and salinity were monitored using data loggers for several weeks at T0, T4, and T12. Cumulative heat stress was estimated by calculating degree heating weeks (DHW) (NOAA Coral Reef Watch 2019; Skirving et al. 2020) using in situ temperature data and the local maximum monthly mean (MMM) temperature of 28.0°C (NOAA Coral Reef Watch station Aruba, Curaçao, and Bonaire). Discrete seawater samples (n=48) were collected opportunistically during environmental monitoring periods to assess nitrate, ammonium, and phosphate at each site. Additional information is provided in the Supplement.

### Phenotypic traits

Partial tissue survival was visually estimated for all fragments at each timepoint using a categorical scoring system. Scores were based on the estimated percentage of living tissue remaining with the following categories: 100, 99-76, 75-51, 50-26, or 25% or less living tissue.

Net calcification rates were determined using the buoyant weight technique (Jokiel et al. 1978). Corals were weighed using a portable balance (precision: 0.1 g, Ohaus Scout Pro, USA) at each timepoint to assess changes in growth across short-term (T0–T4) and longer-term (T0– T12) acclimatization. Daily calcification rates were standardized to surface area (see below).

Dark-adapted respiration (R_dark_), net photosynthesis (P_N_), and light-adapted respiration rates (i.e., light-enhanced dark respiration, LEDR) were measured in the lab using oxygen evolution or consumption in hermetic incubations at each timepoint. PAR and temperature during incubations matched *in situ* conditions measured at the reef and bay site during 7–10 day-long monitoring periods prior to incubations. This allowed us to assess ecologically relevant differences in metabolic rates between transplant groups under realistic temperature and light conditions. Light intensity was maintained using LED lights (Coral Care Gen 1 and Gen 2, Phillips, NL) and relative light spectral profiles were adjusted to maximize similarity to *in situ* conditions (Fig. S3, more info in Supplement). Gross photosynthesis, P_G_, was calculated using the sum of P_N_ and R_dark_. The ratio of daily gross photosynthesis to daily respiration (P:R ratio) was calculated using Equation 1:

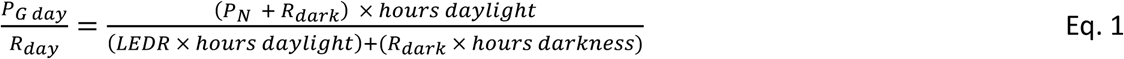

Daylight hours were 12.83 h during T0 and T12 and 11.70 h during T4. The hours of darkness equaled 24 minus daylight hours.

To determine surface area for metabolic and calcification rates, photographs at three angles (0°, 45°, and 90°) were taken full circle around fragments using a turntable and reference ruler (Gutiérrez-Heredia et al. 2015) and rendered into 3D models using photogrammetry software (ReCap Photo, Autodesk, USA). Calibrated models were manually trimmed of non-living area, and live tissue surface areas were extracted using the “mesh report” tool (see Supplement).

For tissue parameters, ash-free dry weight (=tissue biomass) was determined following Fitt et al. (2000) with modifications to use intact pieces of coral tissue and skeleton from an ∼1 cm^2^ piece from the edge or branch tips. Surface area of these pieces was determined using calipers. Chlorophyll a and c^2^ were was extracted in 100% acetone and determined spectrophotometrically using equations from (Jeffrey and Humphrey 1975) and standardized to surface area or number of cells following Schoepf et al. (2015). Symbiont cell density was determined using a Neubauer hemacytometer (*n* = 6 replicate counts per sample). See Supplement for more details on all phenotypic trait measurements.

### Detecting local adaptation and estimating relative fitness

Local adaptation can be assessed using two main criteria to compare the fitness of populations: ‘local vs. foreign’ (fitness within habitats) and ‘home vs. away’ (fitness across habitats) (Kawecki and Ebert 2004). The ‘local vs. foreign’ criterion contrasts the performance of a population within its native habitat relative to foreign transplants whereas the ‘home vs. away’ criterion emphasizes the comparison relative to its own performance in a foreign habitat. While these criteria may be simultaneously satisfied, the ‘local vs. foreign’ criterion is a better diagnostic tool to detect local adaptation driven by divergent natural selection (Kawecki and Ebert 2004). Therefore, from a statistical perspective, detection of local adaptation requires a significant interaction of origin (bay or reef) and treatment (native or transplant), such that native populations outperform foreign populations within the same habitat. In contrast, we refer to environmental specialization when a given population had increased fitness at home vs. away but did not satisfy the local vs. foreign criterion (Kenkel et al. 2015).

The amount of selection (*S*) against transplants was determined using the relative fitness (*W*) of natives and transplants within a given habitat (Hereford 2009). *S* was calculated as follows using the reef-to-bay transplants as an example:

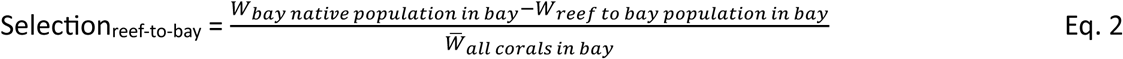

We used calcification rates (T0-T12) and tissue biomass (T12) as proxies for fitness (*W*). Calcification rates have been used as fitness proxies in other work (Kenkel et al. 2015; Castillo et al. 2024) and are an indicator of mass gain as colony size is positively correlated with fecundity (Chornesky and Peters 1987). Tissue biomass was used as a complementary fitness proxy because lipids constitute up to 40% of biomass and are the major component of eggs (Arai et al. 1993), while also promoting increased survivorship during bleaching events (Anthony et al. 2009).

### Statistical analyses

Nonparametric Kruskal-Wallis tests (Scheirer-Ray-Hare extension with *post-hoc* Dunn’s test) were performed to test for differences in (1) daily average and daily variability of temperature, pH_T_, DO, PAR, and salinity, and (2) average inorganic nutrient concentrations between habitats (bay vs. reef) and timepoints (T0, T4, T12).

Linear mixed models (LMM) were used to test for the effects of origin (bay or reef), treatment (native or transplant), and timepoint (T0, T4, T12) on phenotypic traits for each species individually. Random effects of genotype (nested within origin) were included to account for the fact that all parent colonies were equally represented in each treatment as well as repeated measures. For the reaction norms, differences in fitness between the four transplant groups were assessed using the same LMMs as above but without the effect of time. Normality and homoscedasticity of residuals were assessed using Shapiro-Wilk and Levene’s tests, respectively, and Q-Q plots were visually inspected. If assumptions were not met, data was log- or square root-transformed. If assumptions were still violated following transformation, a generalized linear mixed model was performed and models were built using the “family” and link functions that were most appropriate based on the data distribution and Akaike information criterion values. See the Supplement for more details. All statistical analyses were performed in R (v4.2.3) using the “lm4” package (Bates et al. 2015). Results were considered statistically significant at p<0.05.

## Results

### Environmental data

The bay was characterized by more variable and extreme conditions than the reef site with regards to several key environmental parameters, including average daily means and ranges of temperature, pH_T_, DO concentrations, and salinity, though this also depended on the timepoint (Fig. 1E,F,G, Tables S1, S2). Average daily temperatures were significantly higher (1.2–1.3°C) in the bay compared to the reef during T0 and T12, respectively. Daily temperature ranges in the bay were consistently 0.3–0.4°C higher than on the reef, with maximum daily ranges of up to 1.7°C in the bay compared to 1.1°C on the reef. Average DO concentrations were significantly lower in the bay than the reef during T0 and T4 (−4.5% and −11%, respectively) while daily ranges were always significantly higher (+27–286%). Similarly, average pH_T_ was lower in the bay than the reef at T4 (−0.08 pH units) whereas daily ranges also tended to be higher but this was not significant. Average salinity was consistently 5-8% higher in the bay compared to the reef.

Some parameters only showed small or no differences between bay and reef (Tables S1, S2). Average PAR was 11% higher in the bay compared to the reef at all timepoints, but daily maxima were very similar. Nitrate concentrations were unexpectedly higher on the reef (+43%) compared to the bay, regardless of timepoint, while ammonium and phosphate concentrations did not differ between sites or timepoints. Finally, the two reef sites had comparable average daily means and ranges for most abiotic parameters (Tables S1, S2). However, there were a few minor differences at certain timepoints, such as slightly higher daily DO concentrations and pH_T_ or lower temperature range on the transplant reef compared to the source reef.

### Survivorship and partial mortality

A natural heat stress event coincided with the transplant experiment, with cumulative heat stress reaching 18 DHW on the reef during T4 (Fig. 1E). Nevertheless, the large majority of all fragments (95%) survived the experiment, though partial mortality occurred in several transplant groups (Fig. S4 and Supplemental results). Only two fragments had died by T12, one *Porites* reef-to-bay coral (T4) and one *S. siderea* bay-to-reef coral (T12). Transplants from these two groups generally suffered more partial mortality, with ∼10% of *S. siderea* bay-to-reef transplants experiencing ∼26-50% mortality and ∼70% of *Porites* reef-to-bay transplants experiencing ∼50-75% mortality.

### Chlorophyll a and symbiont density

Chlorophyll a concentrations and symbiont densities of *S. siderea* depended on the interaction of timepoint and origin and the main effect of timepoint, respectively, while treatment had no effect. At T4, both reef- and bay-origin corals had lost one-third of their chlorophyll a (−36% and −32%, respectively) and symbiont densities (−38% and −40%) compared to T0, indicating bleaching in response to heat and/or transplantation stress. However, the transplant groups did not show greater bleaching than the native corals in their respective habitat. At T12, both metrics remained significantly lower in all four treatment groups compared to T0, indicating incomplete recovery from heat/transplantation stress, and/or naturally lower levels in late summer/fall independent of origin.

Chlorophyll a and symbiont densities of *Porites* were more affected by transplantation and/or heat stress than *S. siderea,* particularly reef-origin corals (Fig. 2B,D,F). Reef-origin corals lost 54% of their chlorophyll a by T4, while bay-origin corals were unaffected. Reef-to-bay corals, in particular, bleached severely at T4 as they lost 81% of their chlorophyll a and had highly variable symbiont densities compared to reef natives. Bay-to-reef *Porites* exhibited a similar response as reef natives. In contrast to *S. siderea*, both metrics of all four *Porites* groups by T12 were recovered to T0 levels, indicating full recovery from seasonal and/or bleaching-related declines at T4. Similar trends were observed for chlorophyll a per cell, area-normalized chlorophyll c_2_, and chlorophyll c_2_ per cell for both species (Fig. S5; Tables S8, S9).

**Fig. 2.**
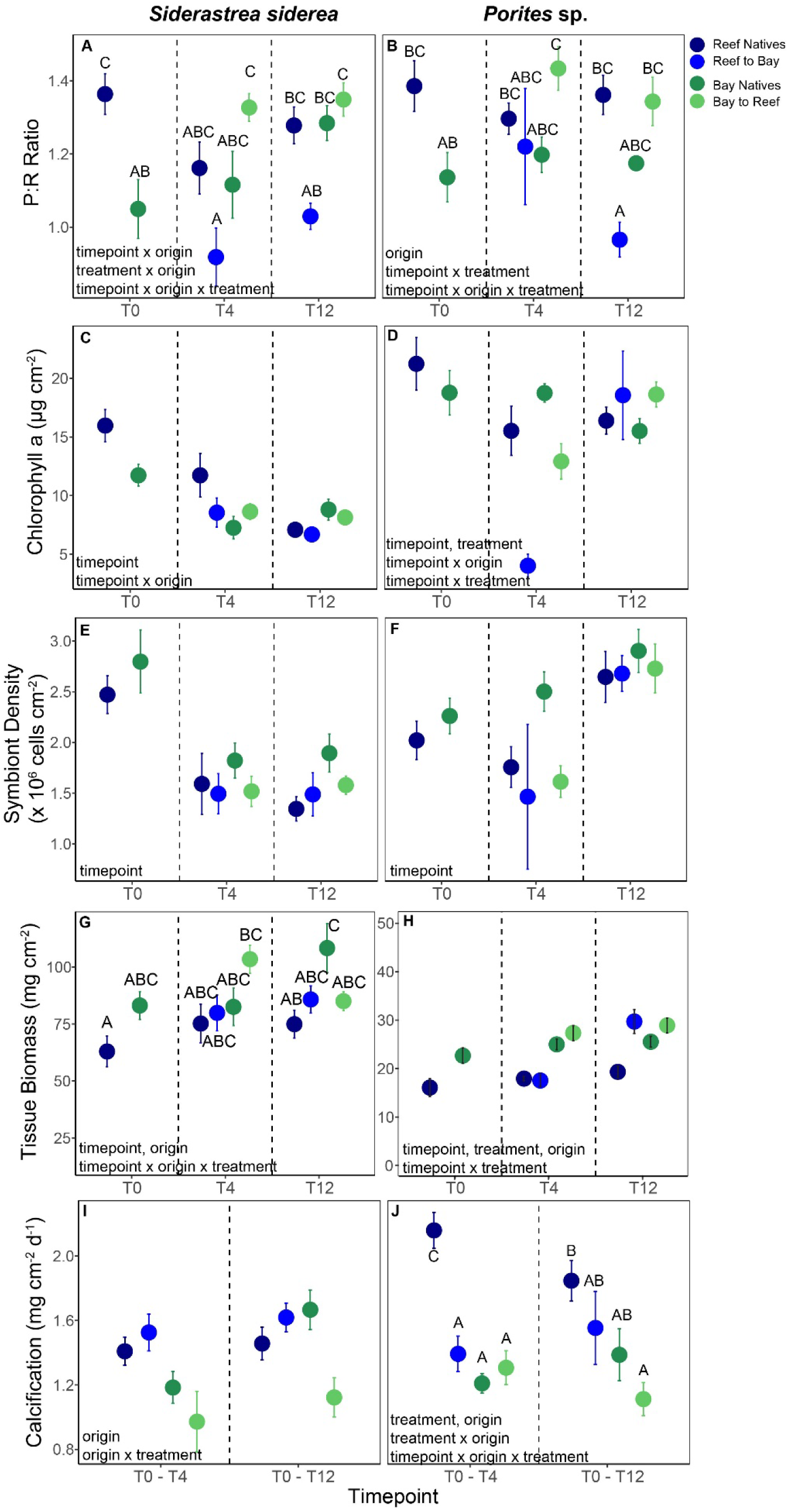
Photosynthesis to respiration (P:R) ratios (A-B), chlorophyll a concentrations (C-D), symbiont densities (E-F), and tissue biomass (G-H) for the four transplant groups of *Siderastrea siderea* and branching *Porites* sp. after 0, 4 and 12 months of transplantation (T0, T4, T12). The y-axes for biomass differ between species. Calcification rates (I-J) are integrated between T0-T4 and T0-T12. Shown is mean ± SE. Significant main effects and interactive terms are indicated in bottom left of each panel. Letters represent *post-hoc* results of pairwise comparisons for three-way interactions only (see Table S5 for all *post-hoc* results)

### Metabolic rates

The P:R ratios of *S. siderea* depended on the interaction of timepoint, origin, and treatment (Fig. 2A, Tables S4, S5). At T0, reef natives had higher P:R ratios (+30%) than bay natives but this was no longer the case at T4, which coincided with significant heat stress (Fig. 1E). The reef-to-bay transplants, in particular, had 18% lower P:R ratios compared to natives at T4, dropping to less than 1, indicating that photosynthesis rates were no longer sufficient to compensate for respiration. After one year, reef-to-bay transplants still had 20% lower P:R ratios (close to 1) compared to both native groups. Reduced P:R ratios of reef-to-bay transplants were due to significantly reduced P_G_ (−39–42%) compared to bay natives, while R_dark_ and LEDR tended to be lower than bay natives, albeit this was not significant (Fig. 2A, Fig. 3A,C,E). In contrast, bay-to-reef transplants were able to maintain similar P:R ratios at T4 and T12 compared to both native groups (Fig. 2A, Tables S4, S5). Bay-to-reef transplants P:R ratios ultimately matched reef natives, however, transplantation resulted in 50% higher P_G_ than reef natives and higher R_dark_ (+44%) and LEDR (+52%), albeit respiration rates were not significantly higher.

**Fig. 3.**
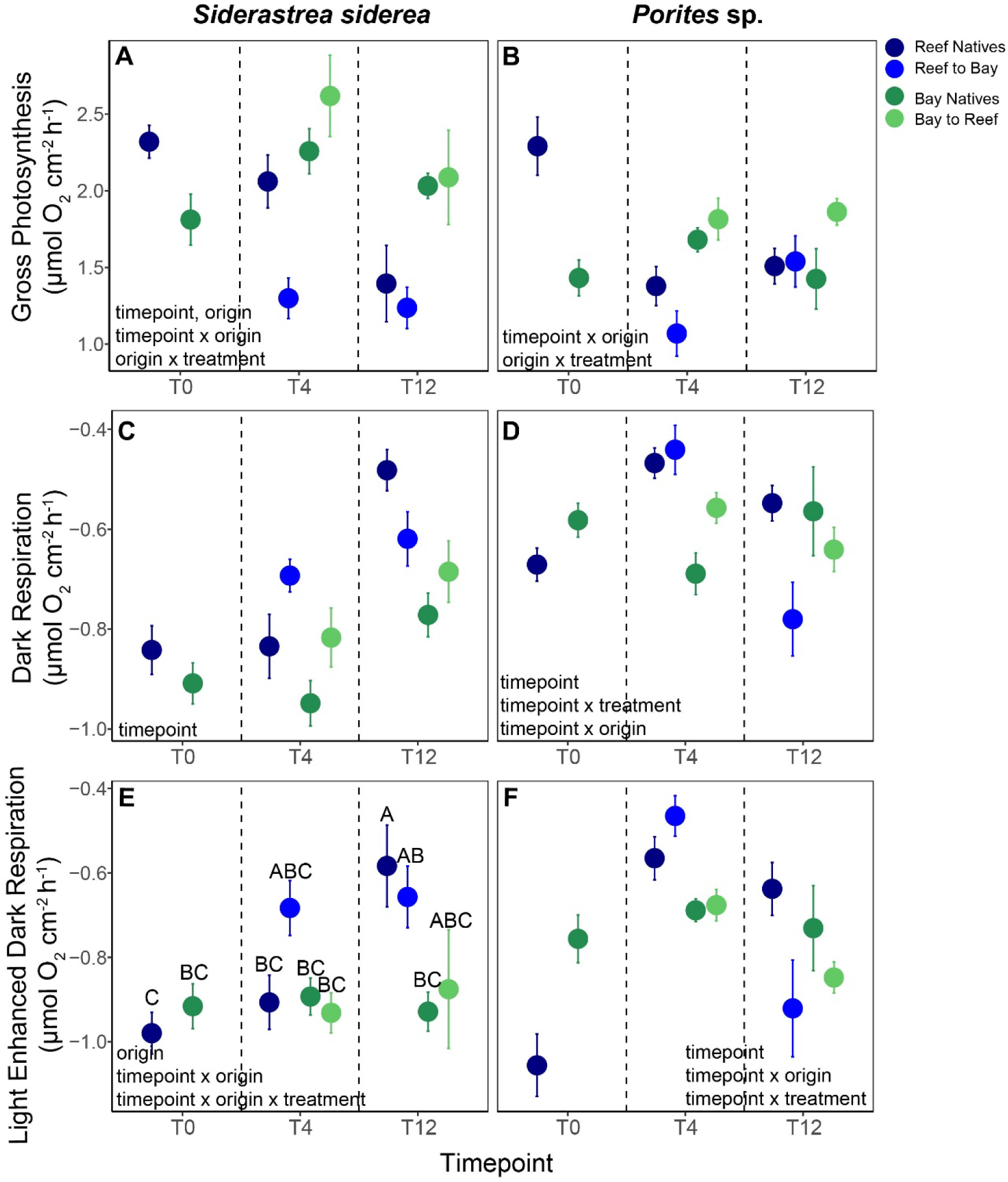
Average rates of gross photosynthesis (A-B), dark respiration (C-D) and light-enhanced dark respiration across all three timepoints (E-F) for the four transplant groups of *Siderastrea siderea* and branching *Porites* sp. after 0, 4 and 12 months of transplantation (T0, T4, T12). Shown is mean ± SE. Significant main effects and interactive terms are indicated in bottom left of each panel. Letters represent *post-hoc* results of pairwise comparisons for three-way interactions only (see Table S5 for all *post-hoc* results)

*Porites* demonstrated similar trends as *S. siderea,* with P:R ratios affected by the interaction of timepoint, origin, and treatment; however, changes in P:R ratios of reef-to-bay transplants were driven by different processes for each timepoint (Fig. 2B, Fig. 3B,D,F; Tables S4, S5). Reef-to-bay transplants had more variable P:R ratios compared to all other groups at T4, but did not show significant declines compared to either natives. By T12, however, their P:R ratios remained 20% lower compared to bay natives and the ratio was <1, indicating potential resource limitation. However, in contrast to *S. siderea*, the reduced P:R ratios of reef-to-bay transplants were primarily driven by significantly higher R_dark_ (+39%) compared to bay natives. Bay-to-reef *Porites* had similar or higher P:R ratios than the reef natives at both timepoints, but had higher P_G_ (+23%), R_dark_ (16%), and LEDR (31%) compared to reef natives.

### Tissue biomass

Tissue biomass of *S. siderea* was affected by the interaction of timepoint, origin, and treatment (Fig. 2G, Tables S4, S5). At T4, reef-to-bay transplants and bay natives had similar tissue biomass, while bay-to-reef transplant had 37% higher biomass than reef natives, albeit not significantly higher. By T12, bay-to-reef transplants’ tissue biomass matched reef natives, while reef-to-bay transplants remained 21% lower than bay natives, albeit not significantly lower.

For *Porites*, bay-origin corals had 25% higher tissue biomass compared to reef-origin corals across all timepoints and treatments (Fig. 2H, Tables S4, S5). Furthermore, there was a significant interaction of timepoint and treatment, with transplants having 31% higher biomass than natives in both habitats but only at T12.

### Calcification

There was a significant effect of origin on *S. siderea* calcification rates, however, this also depended on treatment (Fig. 2I, Tables S4, S5). Reef-to-bay transplants and bay natives had similar calcification rates regardless of timepoint (Fig 2A,I, Tables S4, S5). Conversely, bay-to-reef corals calcified 27% slower than reef natives. Calcification rates were not affected by time (i.e., transplantation duration), remaining similar between T0-T4 and T0-T12.

*Porites* calcification rates depended on the interaction of timepoint, origin, and treatment (Fig. 2J, Tables S4, S5). There was a short-term (T0-T4) effect of transplantation and/or heat stress-related declines in calcification rates of reef-to-bay corals only, which had 34% lower calcification compared to reef natives. However, reef-to-bay transplants still matched the calcification rates of bay natives at T4. In the longer term (T0-T12), calcification rates recovered from transplantation and/or heat stress, leading to intermediate rates of reef-to-bay transplants compared to both reef and bay natives. Bay-to-reef transplants had significantly lower calcification rates (∼40%) compared to reef natives in the short- and long-term.

### Reaction norms

Calcification rates and tissue biomass after 12 months did not provide strong evidence of local adaptation in either bay- or reef-origin corals (Fig. 4, Tables S6, S7). On the reef, native *S. siderea* calcified slightly faster than bay-to-reef transplants, potentially suggesting weak evidence of local adaptation. However, the ‘local vs. foreign’ criterion had insufficient statistical support given that the pairwise comparison was not significant (Fig. 4A). The same pattern was observed for tissue biomass of *S. siderea* in the bay habitat (Fig. 4C). In *Porites*, there was no significant interaction of treatment and origin for calcification rates and the ‘local vs. foreign’ criterion was not satisfied in either habitat for tissue biomass, ruling out local adaptation in both reef and bay populations (Fig. 4B,D, Tables S6, S7).

**Fig. 4.**
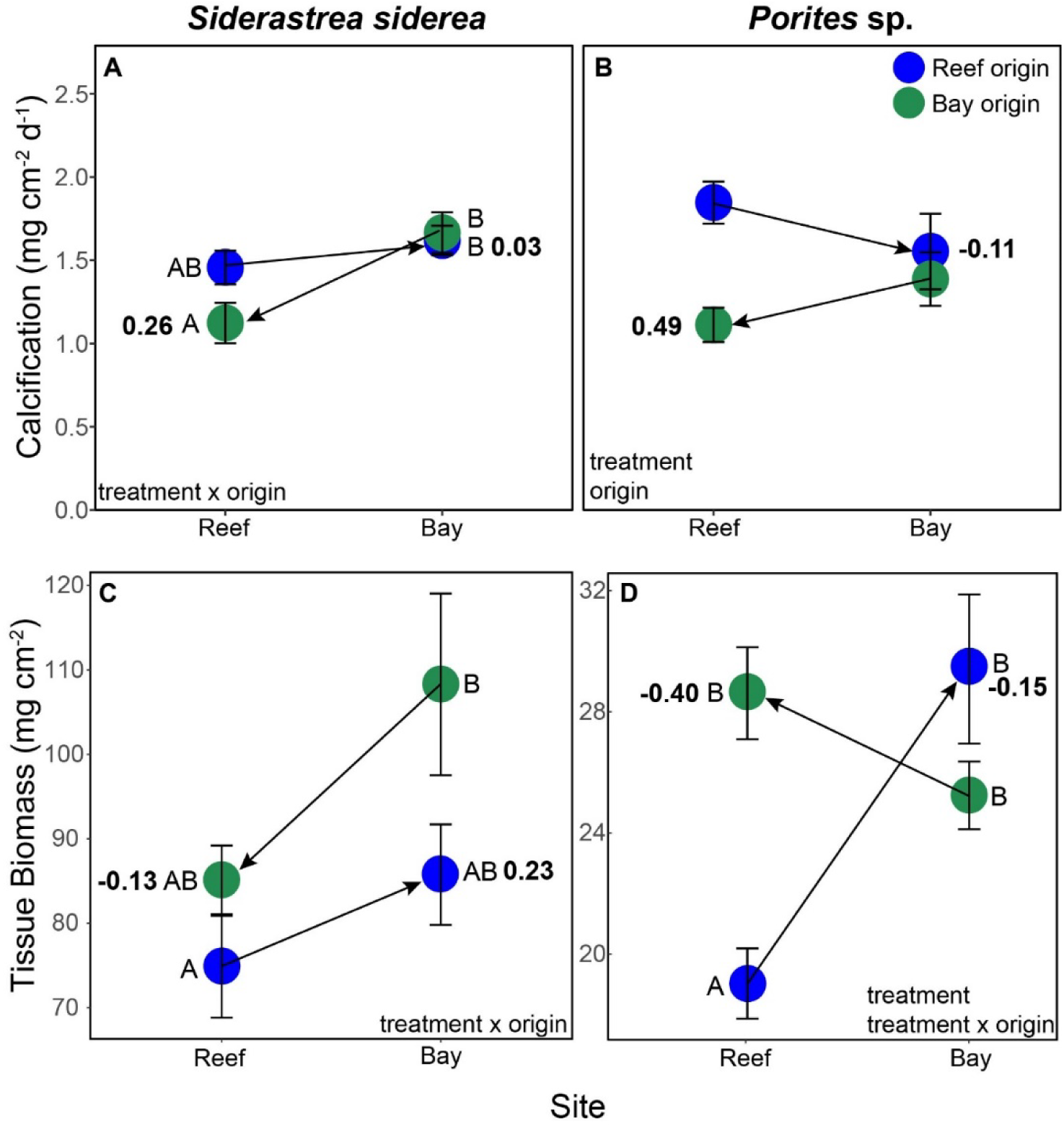
Reaction norms of calcification rates (A-B) and biomass (C-D) of *Siderastrea siderea* and branching *Porites* sp. after 1 year of transplantation. Shown is mean ± SE. Arrows point in the direction of transplantation. Bolded numbers next to transplant groups indicate selection coefficients (*S*), i.e., selection against transplants within a given habitat. Negative *S*-values indicate that transplants have greater fitness than the natives (i.e., maladaptation of natives). Significant main effects and interactive effect of treatment and origin are indicated in bottom left of each panel. Letters represent *post-hoc* results of pairwise comparisons within each panel (see Table S7 for all *post-hoc* results)

Patterns of environmental specialization were observed in both species, but depended on the fitness proxy used and the respective population (Fig. 4, Tables S6, S7). There was no evidence of environmental specialization in *S. siderea* reef-origin corals, as reef natives and reef-to-bay transplants had similar calcification rates and tissue biomass at home vs. away (Fig. 4A,C). Additionally, the selection against reef-to-bay transplants in the bay habitat depended on the fitness proxy used, with calcification rates indicating negligible selection against transplants (*S*_reef-to-bay_ = 0.03) while tissue biomass indicated transplants had a relative fitness disadvantage compared to bay natives (*S*_reef-to-bay_ = 0.23). In contrast, reef-origin *Porites* had 19% higher calcification rates at home vs. away, consistent with environmental specialization (Fig. 4B). However, this pattern was reversed when using tissue biomass as a fitness proxy, where reef-origin *Porites* had 33% lower tissue biomass at home vs. away (Fig. 4D). There was no selection against reef-to-bay transplants using calcification rates or tissue biomass as fitness proxies (*S*_reef-to-bay_ = −0.11 and −0.15, respectively; Fig. 4B,D). Evidence of environmental specialization of bay-origin corals was found for *S. siderea*, as bay-origin corals had 32% higher calcification rates at home vs. away (Fig. 4A, Tables S6,S7). Similar to reef-to-bay transplants, the selection against *S. siderea* and *Porites* bay-to-reef transplants also depended on the fitness proxy used. However, there was a reverse pattern where calcification rates of *S. siderea* and *Porites* indicated relatively more selection against bay-to-reef transplants (*S*_bay-to-reef_ = 0.26 and 0.49, respectively) than tissue biomass (*S*_bay-to-reef_ = −0.13 and −0.40, respectively).

## Discussion

### Acclimatization potential and associated fitness trade-offs in reef-to-bay transplants

We hypothesized that reef-origin corals would follow a specialist strategy and have limited capacity to acclimatize to the more extreme, co-varying multi-stressor bay conditions but would nevertheless survive in the bay, though with significant tradeoffs. Unexpectedly, reef-origin corals from both species demonstrated substantial acclimatization capacity to the bay conditions within one year, as they could adjust all measured phenotypic traits except P:R ratios and tissue biomass to match those of bay natives (Fig. 2,3). However, this plasticity came at the expense of species-specific fitness trade-offs given that P:R ratios and/or tissue biomass of reef-to-bay transplants remained up to 21% lower than in bay natives even after 12 months in their new environment.

The low P:R ratios (∼1) of the reef-to-bay transplants, which were observed in both species (Fig. 2A,B) may have been due to heat-stress induced bleaching, a lack of photo-acclimatization, and/or increased respiratory demand. For *S. siderea*, low P:R ratios at T4 were primarily driven by 42% lower P_G_ rates (Fig. 3A), but transplants had much lower, rather than higher, respiration rates (Fig. 3C,E). At T12, P_G_ remained 39% lower, as did respirations rates. While heat-stress induced bleaching certainly contributed to reduced P_G_ rates, other factors such as incomplete photo-acclimatization may have also played a role. For example, reef-to-bay transplants could have experienced a lag in their photo-acclimatization due to the differing light spectral composition in the bay, which was simulated in respirometry incubations (Fig. S3), or due to changes in their algal symbiont community composition (Little et al. 2004; Cunning et al. 2015).

In contrast, the low P:R ratios of *Porites* reef-to-bay transplants were mostly due to a combination of bleaching (T4) and increased respiratory demand (T12). These corals were much more sensitive to heat stress than *S. siderea* and visibly bleached at T4, as shown by the 81% lower chlorophyll a concentrations relative to bay natives (Fig. 2D). Furthermore, strong declines in both P_G_ and respiration were observed at this time (Fig. 3D). At T12, however, corals had recovered their photo-physiology, but now experienced 44-52% higher respiration rates than bay natives (Fig. 3D) – something that was not observed in *S. siderea*. This may have been a different response to the higher temperatures in the bay (up to 1.3°C warmer during T12), due to increased energetic costs associated with abiotic stress-induced cellular damage, and/or a result of increased heterotrophy (Lesser 1997; Anthony and Fabricius 2000; Clarke and Fraser 2004; Castillo and Helmuth 2005). In other extreme, multi-stress coral habitats, corals also had enhanced respiration rates, potentially indicative of increased heterotrophy (Camp et al. 2017).

The reduced P:R ratios of reef-to-bay transplants of both species (∼20%) reveal a clear energetic cost to living in the bay, but how they impacted resource allocation to skeletal vs. somatic growth, and to what degree they were potentially compensated for by increased heterotrophy, is more equivocal. Interestingly, reef-to-bay transplants maintained calcification rates (T0-T12) in both species (Fig. 2I,J), although corals are typically thought to use autotrophic carbon for calcification and to meet metabolic demand, whereas heterotrophic carbon is mainly used for tissue growth (Hughes et al. 2010). Reef-to-bay *Porites* also had relatively higher biomass than both natives at T12, indicating that heterotrophic feeding may indeed have compensated for the lower P:R ratios (Camp et al. 2017). However, 21% lower tissue biomass in *S. siderea* reef-to-bay transplants (T12) suggests that somatic growth was traded for skeletal growth (Fig. 2G,I). While transplantation did not ultimately result in dramatic reductions of tissue biomass, future work should focus on changes in tissue composition (e.g., lipids, carbohydrates and proteins) and directly measure heterotrophy to help elucidate more fine-scale effects of transplantation on energy budgets and allocation.

### Bay-to-reef transplants have limited acclimatization capacity

Since highly variable, heterogenous environments such as the inland bay (Fig. 1) should favor high phenotypic plasticity and a ‘generalist’ strategy (Kawecki and Ebert 2004), we hypothesized that coral populations from this habitat are able to acclimatize to the more stable conditions on the fringing reef without major fitness trade-offs. Indeed, bay-to-reef transplants of both species adjusted their P:R ratios, chlorophyll a concentrations, and symbiont density to match those of the reef natives after 1 year (Fig. 2A-G). However, they calcified up to 40% less than the reef natives (Fig. 2I,J), and bay-to-reef *S. siderea* additionally exhibited more partial mortality compared to both native groups (Fig. S4A). We therefore reject the hypothesis that bay-origin corals are generalists – rather they showed limited acclimatization capacity consistent with a specialist strategy and experienced fitness trade-offs when exposed to more stable, less extreme environmental conditions, though outcomes may differ when studied over longer time scales (>1 year).

The acclimatization capacity of bay-to-reef transplants stood in stark contrast to that of the other transplant group, with essentially reversed patterns: bay-to-reef transplants adjusted their P:R ratios, but not growth rates, to match that of natives (Fig. 2A,B,I,J). The strong photosynthetic performance under reef conditions was likely mediated by high acclimatization capacity of the algal symbionts or changes in symbiont composition because it is well established that dominant symbiont types often differ across thermally distinct habitats (Howells et al. 2013), though not always (Jung et al. 2021). The reduced calcification rates (27-40% lower) of bay-to-reef transplants in both species may have been driven by the different conditions on the reef such as lower temperature and PAR (Fig. 1, Table S1) (Rodolfo-Metalpa et al. 2008; Ross et al. 2022), however, if environmental control was the primary driver of calcification, then reef natives’ rates should match. Reductions in coral growth were also observed in *Pocillopora eydouxi* when transplanted from a thermally variable to a more stable back-reef pool, while no changes were found for *Porites lobata* (Smith et al. 2008). Combined with findings from other reciprocal transplant studies from nearshore-to-offshore gradients, this highlights that calcification rates can be optimized under more variable temperatures for some species (Smith et al. 2008; Kenkel et al. 2015; Baumann et al. 2021), though other co-varying abiotic drivers may have also played a role. Finally, reduced calcification may also have been driven by a potential shift to more “selfish” symbionts associated with reduced skeletal growth (Little et al. 2004) and reduced transfer of photosynthetic metabolites (Matthews et al. 2018).

It is possible that P_G_ of corals on the reef may have been slightly overestimated because there was a small difference between PAR intensities used in oxygen evolution incubations and long-term averaged PAR intensities measured *in situ* (see Supplementary methods). We based PAR intensities and temperature settings on logger data from a short-term period about ∼10 days prior to incubations at each timepoint. However, PAR levels were ultimately 20-48% higher than the longer-term PAR data recorded for ∼5 weeks per timepoint (Table S1). Nevertheless, we consider it unlikely that this confounds our general findings because bay-to-reef *S. siderea* still had significantly higher (+50%) P_G_ than reef natives when incubated under identical PAR intensities, and a similar trend (+23%) was observed for *Porites* (Fig. 3A,B).

### The role of environmental specialization in shaping acclimatization capacity

Using calcification rates and tissue biomass as proxies for coral fitness, we found no compelling evidence of local adaptation of either reef or bay *S. siderea* or *Porites*. Although significant origin x treatment effects were observed for all of these fitness proxies after 1 year of transplantation except calcification in *Porites*, the ‘local vs. foreign’ criterion was not met in either population (Fig. 4A-D). Reef-origin corals further showed little evidence of environmental specialization based on the ‘home vs. away’ criterion (Kenkel et al. 2015), as fitness proxies were typically lower at home vs. away, again with the exception of calcification rates in reef *Porites*. This is in contrast to work from Belize where forereef-sourced *S. siderea* were more environmentally specialized than nearshore and backreef conspecifics (Castillo et al. 2024), or other studies demonstrating local adaptation in corals from cooler, more stable reef habitats (Kenkel et al. 2013; Rivera et al. 2022). Overall, reef-origin corals of both species therefore seem to follow a ‘generalist’ strategy with substantial acclimatization capacity (albeit fitness trade-offs), which is consistent with the largely low selection pressure against reef-to-bay transplants (i.e., low *S*-values) and their overall plastic physiological response during the experiment (Fig. 2,3).

Compared to reef-origin corals, bay-origin corals of both species showed greater evidence of environmental specialization, but this also depended on the fitness proxy used. Calcification rates supported specialization more than biomass, as also indicated by greater selection against bay-to-reef transplants (Fig. 4A,B). Bay *S. siderea* generally showed stronger evidence of environmental specialization than bay *Porites* because they had significantly higher (32%) calcification rates at home vs. away while biomass also tended to be higher (Fig. 4A,C). Environmental specialization of calcification rates has also been detected in nearshore *S. siderea and Pseudodiploria strigosa* from Belize and *P. astreoides* from the Florida Keys (Kenkel et al. 2015; Baumann et al. 2021). In contrast to *S. siderea*, calcification rates of bay *Porites* were somewhat higher at home vs. away but no treatment x origin interaction was detected, and biomass was similar across both habitats (Fig. 4B,D); thus, they mostly follow a generalist strategy similar to the reef-origin corals.

Environmental specialization of inland bay-origin corals in this study and other Caribbean nearshore species (Kenkel et al. 2015; Baumann et al. 2021) is counterintuitive to theory that suggests exposure to more heterogeneous physicochemical conditions should promote phenotypic plasticity (Kawecki and Ebert 2004). One possible explanation could be that environmental variability in the bay is high in terms of amplitude, but predictability is low, thus leading to reduced phenotypic plasticity (Bitter et al. 2021). Lower calcification rates under more homogeneous, stable conditions could also represent the costs of possessing phenotypic plasticity in other traits, such as tissue biomass (Kenkel and Matz 2016; Rivera et al. 2021). For example, nearshore *P. astreoides* had higher calcification rates at home vs. away, however, they nonetheless had higher baseline capacity to match their gene expression profiles to those of offshore natives compared to offshore-to-nearshore corals (Kenkel and Matz 2016). Yet, environmental specialization of nearshore corals is not ubiquitous, for example, nearshore *S. siderea* in Belize had similar calcification rates across all habitat types following 3.5 years of transplantation to nearshore, backreef, or forereef habitats (Castillo et al. 2024). Given contrasting patterns of environmental specialization across similar environmental gradients (i.e., more stable to more variable habitats) found in other studies (Kenkel and Matz 2016; Baumann et al. 2021; Castillo et al. 2024), more work is needed to fully understand how multiple co-varying stressors shape corals’ adaptive capacity, how this is influenced by environmental predictability, and which phenotypic traits best predict fitness.

The initially unplanned switch from Tugboat Reef to Carmabi House Reef as the reef transplantation site became necessary due to sudden partial mortality observed at Tugboat Reef during a separate pilot transplantation study (see Supplement for details). However, in-depth abiotic monitoring showed that mean temperature, pH, DO, salinity, PAR, and nutrients were highly similar between the two reef sites at all three timepoints, and different between both reef sites and the bay (Fig. 1E, Table S1). Any physiological changes in the reef natives between T0 and the later timepoints can also reflect seasonal changes in physicochemical conditions and/or heat stress during T4. Thus, while it is possible that they initially experienced some transplant stress, we are confident that our main findings for this 1-year experiment are valid.

## Conclusions

Here, we demonstrate that two coral species with different life history strategies have substantial phenotypic plasticity to acclimatize to novel highly variable, multi-stressor conditions within one year, providing crucial insights into coral adaptive capacity under intensifying global and local stressors. This finding provides hope for the persistence of coral reefs under future ocean conditions, though the compromised P:R ratios of reef-to-bay transplants indicate fitness trade-offs that require compensation strategies such as increased heterotrophy for long-term success and survival. It is further likely that the 1-year exposure to the highly variable conditions of the bay has led to increased stress tolerance of the reef-to-bay transplants (Safaie et al. 2018; Brown et al. 2022), particularly with respect to their heat tolerance, given that the bay is not just thermally more variable, but also ∼1°C warmer than the reef (de Jong et al. 2025). Future work should therefore compare the resistance of bay- and reef-origin corals to heat and other climate change stressors and also address the role of the coral microbiome and potential host cryptic speciation in modulating coral adaptive capacity (Grupstra et al. 2024; Aichelman et al. 2025)

Corals originating from the bay also had substantial capacity to acclimatize to more benign, stable environmental conditions on the reef, but their growth may be only optimized in their native habitat due to environmental specialization. More direct measures of fitness (e.g., fecundity) and genetic analyses are needed to confirm this result. This finding is consistent with other work from the Caribbean, though from less extreme environmental gradients (Kenkel et al. 2015; Baumann et al. 2021; Castillo et al. 2024). Given their environmental specialization, bay-origin corals may not be the most suitable broodstock to supply naturally stress-resistant corals for proactive restoration approaches, though some fitness trade-offs may become acceptable under intensifying climate change (Caruso et al. 2021). Overall, our study highlights the important role that natural laboratories with extreme environmental conditions play in investigating coral adaptive capacity to multiple, co-varying stressors under ecologically realistic conditions (Camp et al. 2017; Schoepf et al. 2023).

## Acknowledgements

We thank M.J.A. Vermeij, A. Dulskiy, P. Hernandez, and the staff at CARMABI Research Station for their assistance in the field; V. Chamberland, K. Latijnhower, M. Achlatis, and J.D. Goeij for the use of their equipment in the field; I. Benda, I. Guncay, B. van Beusenkom, P. Slot, and M. Schuurmans for laboratory technical assistance.

## Author Contribution Statement

SLS: Investigation, Formal Analysis, Visualization, Writing – Original Draft Preparation, Funding acquisition

CRAL: Investigation, Formal Analysis, Writing – Review & Editing

RM: Investigation, Formal Analysis, Visualization, Writing – Review & Editing

MEP: Investigation, Visualization, Writing – Review & Editing

KWJ: Investigation, Writing – Review & Editing

SS: Investigation, Formal Analysis, Writing – Review & Editing

RMZ: Formal Analysis, Writing – Review & Editing

VS: Conceptualization, Investigation, Writing – Review & Editing, Funding Acquisition, Resources, Project Administration, Supervision (lead)

## Statements and Declarations

### Ethics

Research was conducted under permit #2019/021824 issued to the CARMABI Foundation by the Curaçaoan Ministry of Health, Environment and Nature.

### Data availability statement

All data and code will be provided following publication on figshare.com.

