## Supplemental Information for "Physiological mechanisms underlying coral acclimatization capacity to novel, multi-stressor conditions"

Title:

Sarah L Solomon^1^ 
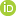
 ^*^, Christian RA Lippens^1^, Riccardo Mazza^1^, Maya E Powell ^2^, Kelly W Johnson^1^, Sophia Suvacarov^1^, Rene M van der Zande^1^, Verena Schoepf^1^

^1^Department of Freshwater and Marine Ecology, Institute for Biodiversity and Ecosystem Dynamics, University of Amsterdam, Amsterdam, the Netherlands

^2^Environment, Ecology, and Energy Program, University of North Carolina at Chapel Hill, Chapel Hill, NC, 27514, USA

*Corresponding author:

Sarah L Solomon;

**Supplemental Methods**

**Study sites, coral collection, and experimental design**

**Spanish Water Bay description**

The physico-chemical conditions and water quality of Spanish Water Bay and fringing reefs on Curaçao are described in detail elsewhere (de Jong et al. 2025). The semi-enclosed geomorphology, the longer seawater residence times, and relatively shallow depth (5-12 m) of the bay lead to 0.3-0.8°C higher average temperatures depending on the season and relatively higher daily variability of temperatures in the bay (maximum daily range: 1.3-1.6°C) compared to a reference reef site, Spanish Water Reef (0.6-0.8°C) (de Jong et al. 2025). The bay is surrounded by mangroves and the benthic substrate consists of mud, soft, fine-grained sediment, sand, and rubble. There are currently no framework-building reefs in the inland bays, rather they are dominated by seagrass, macro- and turf algae interspersed with isolated coral assemblages (Fig. S1, Kuenen and Debrot 1995). The benthic substrate and/or community contribute to naturally high turbidity, lower light levels, and highly variable pH (maximum daily range: 0.13-0.25 pH units, range: 0.33 pH units) and dissolved oxygen concentrations (maximum daily range: 3.15-5.91 mg L^-1^, range: 6.45 mg L^-1^) (de Jong et al. 2025). Nutrient input via terrestrial run-off contributes to elevated nitrate, ammonium, and phosphate concentrations in the bay compared to the reference reef site, especially during the wet season (Kuenen and Debrot 1995; de Jong et al. 2025). Further, effect-based chemical water quality assessment has revealed that organisms in the bay are exposed to higher ecotoxicological risks than organisms at the reference reef site through chemical pollution (de Jong et al. 2025).

Note that our study site in Spanish Water Bay was located around the corner from the Windsurfing Club ‘Windsurfing Curaçao’ (12°04'36.4" N, 68°51'35.8" W) whereas the monitoring site in (de Jong et al. 2025) is located ~500 m away (12°04′15.35” N, 68°51′30.61” W). Similarly, the reef site, Tugboat Reef, (12°04'04.9" N, 68°51'42.7" W) is located ~350 m away from the reef monitoring site, referred to as Spanish Water Reef or Director’s Bay, (12°03′53.48” N, 68°51′36.69” W) in (de Jong et al. 2025).

***Differentiation between branching Porites* spp*.***

The taxonomic status of branching *Porites* species in the Caribbean (i.e., *P. divaricata*, *P. furcata*, and *P. porites*) has been a topic of debate due to high morphological plasticity, with some studies finding near-continuous morphological variation between the three species (Prada et al. 2014). This made species ID during field collection challenging. In this study, we aimed to collect corals with as much morphological similarities as possible within and between bay and reef habitats. We aimed to collect *P. furcata* that were characterized by skinny branches, bifurcation closer to the base of colonies, and shorter polyps relative to what is usually described for *P. porites* (Fig. S1).

**Coral collection, fragmentation, and physical structures of transplant experiment**

To minimize the probability that identical genotypes (= genets) were collected, colonies of the same species were selected at least 5 m apart from one another. Intact colonies were transported to Carmabi Research Station in shaded and aerated plastic tubs filled with seawater. On the same day of collection, each of the parent colonies were cut into five fragments (= ramets) of roughly equal size using a bandsaw (~6 cm diameter for *S. siderea* and one ~6 cm-length branch for *Porites*). One fragment from each colony was immediately frozen at -80 °C for baseline (T0) measurements of tissue parameters, while the other 4 fragments were buoyant weighed (see below) and glued to labeled acrylic tiles using thick viscosity cyanoacrylate adhesive (Glue Masters LLC). Acrylic tiles with fragments were fastened to larger acrylic plates that allowed for transportation submerged in seawater-filled tubs between research station and study sites. Plates with corals were then affixed to PVC structures that were supported by rebar such that the acrylic plates were raised ~20 cm from the seabed. Tiles and plates were cleaned every 1.5 months. Prior to transplantation, corals were kept at temporary holding sites near Carmabi Research Station for 15 days (holding site locations: 12°07'20.7"N 68°58'10.3"W for reef-origin corals and 12°07'30.5"N 68°58'05.7"W for bay-origin corals). During these 15 days, plates with corals were brought back into the lab every other day to measure surface area and metabolic rates. All corals were given 5-6 days to recover from collection/fragmentation prior to measurement of metabolic rates. After 4 months of transplantation (=T4, June 2022 – October 2022), half of the corals from each transplant group were collected for live measurements and subsequently frozen at -80°C. The remaining fragments stayed in their native or foreign habitat until 12 months post-transplantation (=T12, June 2022 – June 2023), when they were collected for live measurements and subsequent transport in a dry shipper and storage at the University of Amsterdam at -80°C for further tissue analysis.

**Change in reef transplantation site**

Reef corals were collected from Tugboat Reef. Following fragmentation and attachment to labeled tiles, reef-origin corals were allowed to recover for 15 d on a temporary holding structure on Carmabi House Reef. During this time, a different set of corals from a separate pilot transplantation study (also collected from Tugboat Reef and Spanish Water Bay) were transplanted back to Tugboat Reef for 7 d. Unexpectedly, these corals experienced extensive partial tissue mortality, which may have been due to predation and/or a strong stress response. The upper ridges of *S. siderea* corallites were drastically white, while polyps remained unbleached. Tissue was slowly sloughing off the skeleton, which suggested a stress response that was leading to tissue necrosis (Fig. S2). We observed this response on small patches along *Porites* *furcata* branches as well, but to a lesser extent. Although the exact cause of mortality and the stress response remains unknown, we suspect that this was the result of microbial infection of an opportunistic pathogen on wounded edges of fragmented *S. siderea* colonies and/or predation by *Stegastes planifrons*, an algae-farming damselfish, or the fireworm *Hermodice carunculata*, a known corallivore (Wolf et al. 2014; Vermeij et al. 2015). Large numbers of this damselfish and the fireworms were observed at Tugboat Reef at this time. We therefore decided to put the pilot study corals back to the holding site at Carmabi House Reef where their health condition improved rapidly. These pilot study corals were not used for the transplant experiment described here. No stress response was observed in the corals used for the transplantation experiment described here during their recovery from fragmentation at Carmabi House Reef. This shows that the stress response of the pilot corals was likely not the result of handling or fragmentation, but was specific to a stressor at Tugboat Reef at that time. To avoid coral loss that would be unrelated to transplantation and put the transplant experiment at risk, we therefore chose Carmabi House Reef as the replacement reef site due to its similarities to Tugboat Reef, including the relatively stable and similar environmental conditions, and closeness to another inland bay (Piscadera Bay). Reef natives and bay-to-reef transplants thus stayed at Carmabi House Reef for the entire 1-year duration of the transplant experiment.

**Environmental monitoring**

Abiotic data was monitored using in situ data loggers (logging intervals of every 10 min) attached to PVC coral transplantation structures at the bay and the two reef sites. Temperature was continuously monitored over the course of the entire experiment using Envloggers (T7.3, ElectricBlue, PT, ≤ 0.1°C precision and ≤ 0.2°C accuracy). Envloggers were not wrapped in tape or otherwise shaded which potentially introduced temperature bias under contrasting irradiances (Rich et al. 2024). However, the average PAR levels at the sites in our study were, on average, equal (Table S1) and significant bio-fouling over the 1-year deployment period would have further minimized solar heating bias. Note though that Envloggers may generally overestimate the true temperature by ∼0.2°C (Rich et al. 2024). Although other loggers also recorded temperature (see below), only temperature data from the Envloggers were used to characterize thermal regimes and calculate cumulative heat stress. PAR was semi-continuously monitored between 28 May - 26 August 2022, 5 October – 13 November 2022, and 18 May 2023 – 12 June 2024 whereas pH_T_, DO,and salinity were monitored between 28 May – 24 June 2022, 5 October – 13 November 2022, and 18 May 2023 – 12 June 2023. PAR loggers (Xtreem Photosynthetic Active Radiation Logger, Odyssey, NZ) were calibrated via simultaneous deployment with a miniPAR logger (PME, Inc, USA, up-to-date factory calibration; accuracy of ± 5% in air) that were all set to 1 min logging interval during calibration. A calibration coefficient was calculated by linear regression and applied to data from each Xtreem Photosynthetic Active Radiation logger. pH loggers (HOBO pH and temperature logger MX2501, Onset Brands, USA; accuracy of ±0.10 pH units) were equipped with copper guards to prevent biofouling. pH loggers were retrieved and re-calibrated approximately every other day using TRIS buffer purchased from A.G. Dickson (Scripps Institution of Oceanography, USA) to record pH on the total scale and then re-deployed. The calibration was performed by measuring the mV of TRIS buffer using a Hach multimeter (HQ40d, Hach, NL) at two temperatures within the range of ambient seawater temperatures (i.e., 26-30°C) in order to determine seawater pH on the total scale following (Dickson et al. 2007). pH data from up to 24 hours before and after calibration were considered reliable due to known drift of glass pH sensor probes. DO loggers (miniDOT, PME, Inc, USA; accuracy of ± 5% of reading or ± 0.3 mg L^-1^) were factory calibrated prior to use and sensor tips equipped with either a copper ring (at bay sites) or copper mesh (at reef sites) to prevent biofouling. The factory calibration was verified by submerging DO loggers into 100% air saturated seawater (aerated with air pump) and into 0% saturated seawater (aerated with N_2_ and/or adding sodium sulphite) and cross-checking measurements with a calibrated Hach multimeter. Conductivity loggers (Conductivity and Temperature, Odyssey, NZ; accuracy within 3% of reading) were calibrated by applying unique calibration equations provided by the factory and calibration was verified by submerging loggers into an aqueous solution of 35 ppt NaCl. Conductivity loggers were mounted at least 10 cm away from any objects. Conductivity (mS cm^-1^) was converted to salinity using the marelac package in R (Soetaert et al. 2023) using the conductivity ratio (= conductivity divided by standard conductivity of seawater at salinity = 35 ppt, temperature = 15°C, and pressure = 0 bar), temperature, and pressure (*P*) against the local atmospheric pressure. *P* (in Pascal) was determined using the following formula: *P* = $\rho$ * *g* * *h*; where *ρ* = density of seawater at given temperature, *g* = gravity velocity = 9.81 m s^-2^, and h = depth (m). The average seawater temperature measured by the conductivity loggers during each timepoint was used to determine $\rho$. DO data was adjusted according to matching salinity data following methods from (Garcia and Gordon 1992). All loggers were cleaned every other day during monitoring periods.

Cumulative heat stress was estimated by calculating degree heating weeks (DHW) (NOAA Coral Reef Watch 2019; Skirving et al. 2020) using in situ temperature data and the local maximum monthly mean (MMM) temperature of 28.0°C (NOAA Coral Reef Watch virtual station Aruba, Curaçao, and Bonaire). We only calculated cumulative heat stress for the reef habitat (Carmabi House Reef site) because historical temperature data are not available for the bay and the spatial resolution of the Coral Reef Watch satellite data is too coarse to distinguish between reef and bay habitats. Furthermore, given that the bay is on average 0.8-1.3°C warmer than the fringing reef (de Jong et al. 2025), it is also likely that bay corals experience higher long-term MMM temperatures and, thus, have higher bleaching thresholds. Therefore, we do not consider the local Coral Reef Watch bleaching threshold (~29.0°C) to be applicable to the bay-origin corals and did not calculate separate DHW for the bay site.

***Inorganic nutrients***

Discrete seawater samples were opportunistically collected during all three environmental monitoring periods to measure inorganic nutrients concentrations (ammonium, nitrate, phosphate). Water was collected at each transplant site in plastic bottles and immediately filtered (0.2 μm polyethersulfone filter, VWR, NL). Water samples for nutrient analyses (n = 48) were filtered into vials (7 mL Pony Vials™, PerkinElmer, NL), transported in a cooler on ice, and stored at -20 °C until analysis of nitrate, nitrite, ammonium, and phosphate using a San ++ Automated Wet Chemistry Analyzer (Skalar, NL).

**Coral phenotypic traits**

**Calcification rates**

Coral fragments and tiles were initially weighed after fragmentation by suspension in seawater using a plastic platform and string connected to a portable balance. After corals were glued to their tiles, the fragments were weighed again. At T4 and T12, tiles with fragments were carefully cleaned and weighed again. Using equation (2) derived from Jokiel et al. (1978), we determined the dry weight of each fragment using the density of seawater (calculated based off temperature and salinity at time of measuring) and aragonite (2.93 g cm^-3^). Daily area-normalized calcification rates were obtained by calculating the difference in dry weight between T0 and T4 and T0 and T12, and dividing this difference by the number of days elapsed. Average surface area (from photogrammetry measurements) between each set of timepoints was used to standardize calcification rates to tissue surface area (Foster et al. 2014).

**Metabolic rates**

Corals were retrieved from their sites one day prior to incubations and dark-adapted in temperature-controlled indoor aquaria at the Carmabi Research station overnight. Between 9:00 and 16:00 the next day, each fragment was placed on a stainless steel stand in a clear acrylic chamber that was filled with filtered seawater (20 µm), fit with a stir bar, and a calibrated fiber optic oxygen sensor (OXY-4 SMA and mini, PreSens, DE). Chambers were semi-submerged in temperature-controlled aquaria with heaters (ThermoControl 200, EHEIM GmbH, DE) and aquaria pumps (Nano Voyager 1000 L h^-1^, Sicce, IT). Rates were converted from % air saturation min^-1^ to O_2_ mg h^-1^ by using the solubility of oxygen at the incubation temperature and salinity, and by normalizing the values to the volume of seawater in each chamber. Rates were standardized to each fragment’s surface area (see below). Rates were determined from a standardized selection of the slope of each line of % air saturation over time and background signal was subtracted using blanks from each trial or daily average of blanks. Light intensity in incubations was measured in the aquaria using a submersible PAR sensor (MQ-510 Quantum Sensor, Apogee, USA).

***Incubation protocol modifications between habitats and timepoints***

Incubation conditions for the initial timepoint (T0) were based off of monitoring at the Spanish Water bay and reef sites in previous years (de Jong et al. 2025). At T4 and T12, we measured temperature and light at our bay and reef site over a 7 – 10 d monitoring period prior to incubations, and adjusted these abiotic conditions in our aquaria and incubations accordingly (Table S2). Spectral light data at coral collection depths were available for Spanish Water Bay (Fig. S3A) and a reef site in Curaçao that has a presumed similar light environment to Tugboat Reef (Santa Martha Reef, 12°15’59.56”N, 69°07’39.66”W; Fig. S3B). These spectral measurements were made in November 2021 using a hyperspectral radiance sensor (RAMSES, TriOS Mess- und Datentechnik GmbH, DE). Since the light spectrum in the bay was characterized by higher intensities of green, yellow, and red light (550 – 665 nm), we adjusted the spectra of the Phillips CoralCare Gen 1 and Gen 2 used for aquaria and incubations as closely as possible to *in situ* spectra using the “% warm” and “% cool” settings in the CoralCare software (Fig. S3). Corals from the bay transplant site were incubated under 100% warm light and reef transplant site corals were incubated at 50% warm and 50% cool (Fig. S3C).

Some modifications were made to respirometry incubation protocols following T0. Namely, respirometry incubations were, on average, 30 min shorter at T4 and T12. At T0, incubations were only terminated after a 5% change in dissolved oxygen saturation relative to the blanks, while at T4 and T12, incubations were terminated after about a 2% change because a representative slope was already observed after a 2% change. Additionally, at T0, dark respiration was measured on a different day than net photosynthesis and light-enhanced dark respiration whereas at T4 and T12, we measured dark respiration, net photosynthesis, and light-enhanced dark respiration consecutively.

**Surface area measurements using 3D photogrammetry models**

Two sets of photographs were taken full circle around coral fragments using a turntable and high contrast reference ruler, as described in (Gutiérrez-Heredia et al. 2015). Briefly, the camera (LUMIX DMC-TZ70, Panasonic, Japan) lens was placed level with the coral fragment, thus creating photos from a ground (0°) angle. The turntable was turned ≈15° after every photo, resulting in approximately 20-25 successive, overlapping photos per coral at this angle. For the second set of photos, the camera was placed at a ≈45° angle and a similar pattern of photos was taken, turning the turntable ≈15° after every photo, resulting in another 20-25 photos. Finally, the camera was held directly above the coral (90°) and a single calibration photo was taken, ensuring the ruler was fully in frame. The entire process thus yielded a total of 50-60 photos per coral and took no more than approximately 3 min per fragment, ensuring minimal air exposure of corals. All photographs were taken outdoors under cover to avoid direct light. Photos were uploaded to Autodesk ReCap Photo software (versions 1850 – 1740), which rendered photos into 3D models. Each model was calibrated from the top view using the “scale by” function by selecting two in-focus points on the reference ruler and entering the distance between points. Using “window select” and “brush” tools, all areas of the models excluding living coral tissue were deleted. The surface area and volume of each coral fragment were recorded using the “mesh report”.

**Tissue biomass**

For *Siderastrea siderea*, a rectangular piece of tissue and skeleton was cut from the edge of fragments using a hand saw. Surface area was determined by assuming a rectangular shape and measuring the width and height of each piece using calipers. For *Porites*, a branch tip was used and surface area was calculated by assuming a cylindrical shape and measuring the radius and height of each tip. Tissue and skeleton were dried at 60°C for 72 h in pre-combusted aluminum pans and weighed on an analytical balance (ABT 220-5DNM, Kern & Sohn, Germany; linearity: ± 0.0001). Following combustion at 450°C for 6 h, pans were weighed again. The ash-free dry weight was calculated as the difference between dry and burnt weight and standardized to surface area.

**Chlorophyll concentrations and symbiont cell density**

Coral tissue was removed from the skeleton using a WaterPik flosser (waterpik®, USA) filled with ultrapure MilliQ Type 1 water (MilliQ®, Millipore, Merck KGaA, DE) and a 1 cm^2^ stencil. The tissue slurry was homogenized for 30 sec (Tissue-Tearor™, Model 985370, BioSpec Products, Inc, USA) and centrifuged at 1500 g for 10 min at 4°C to pellet the algal cells. After the supernatant was removed, the algal pellet was rinsed with 2 mL of MilliQ, centrifuged a second time at 1500 g for 10 min, and the supernatant was removed. The pellet was resuspended in 5 mL of MilliQ and 2 – 3 mL were aliquoted for chlorophyll chl a + c^2^ concentrations and 1 mL was aliquoted for symbiont cell density. Samples were stored at -20°C until further processing. Algal aliquots for chlorophyll extractions were centrifuged at 3500 g for 10 min at 4°C and the supernatant was removed. Chlorophyll was extracted in the dark at -20 °C for 24 h using 100 % acetone. After re-centrifuging, area-normalized chlorophyll a and c^2^ concentrations were determined spectrophotometrically (Novaspec III, Amersham Biosciences, UK) by measuring absorbance at 630 and 663 nm, respectively. Additionally, absorbance was measured at 750 nm to estimate sample turbidity and this value was subtracted from the absorbances at 630 and 663 nm. Chlorophyll concentrations were calculated using equations from (Jeffrey and Humphrey 1975), and standardized to surface area. Symbiont cell densities were determined by counting 6 replicate aliquots of 0.1 µL on an improved Neubauer hemacytometer, and standardized to surface area. Surface areas were pre-defined with a stencil during tissue removal.

**Statistical analyses**

For LMMs, the significance of fixed effects was determined using Type-III sum of squares and the Satterthwaite’s method for approximating degrees of freedom using the lmerTest package (Kuznetsova et al. 2017). The significance of the random effect of genotype was evaluated using likelihood ratio tests (LRT) after reducing or removing the random-effect term using the ranova() function from the lmerTest package (Kuznetsova et al. 2017). Main and interactive effects were considered significant at p<0.05. Pairwise contrasts were generated using the emmeans package (Lenth et al. 2025) when main and/or interactive terms were significant, with p-values adjusted using the Tukey method and the Kenward-Roger degrees-of-freedom method. Further, compact letter display of all pairwise comparisons were generated using the cld() function from the multcomp package (Piepho 2004; Hothorn et al. 2008) where different letters indicate a significantly different pairwise contrast. For GLMMs, the significance of fixed effects was determined using Analysis of Deviance tables with Type III Wald chisquare tests using the Anova() function from the car package (Fox and Weisberg 2019). GLMMs were built using the Gamma distribution with the “identity” (no link) was used in all cases except in case of R_dark_ *S. siderea*, where the “log” link function was used. The absolute value of respiration rates was used as the Gamma distribution is only defined for positive values. In the event of failed convergence of GLMMs, models were re-fit with a range of optimizers using the allFit() function in the lme4 package and in all cases, the Bound Optimization BY Quadratic Approximation, or “bobyqa”, optimizer was used because models fit with this optimizer converged the fastest.

**Supplemental Results**

**Survivorship and partial mortality**

*Siderastrea siderea* reef natives, bay natives, and reef-to-bay transplants experienced the least partiality mortality, with 20-30% of fragments suffering a maximum of ~25% partial mortality by T12 (Fig. S4). *Siderastrea siderea* bay-to-reef transplants had higher partial mortality than the other three groups, with 44% of fragments experiencing ~25% mortality and 11% experiencing ~50% mortality. Reef and bay native *Porites* also suffered a maximum of ~25% partial mortality, with more bay native fragments (63%) experiencing partial mortality compared to reef natives (20%). Reef-to-bay *Porites* had the highest rates of partial morality of any group, with 72% of fragments experiencing either ~50 or ~75% mortality. Bay-to-reef transplants had similar partial mortality rates as reef natives, with 30% of fragments experiencing ~25% mortality and 10% of fragments experiencing ~50% mortality. Some of the partial mortality observed in corals at the bay site was partially due to parasitization by an unknown species of clams at T12 only. This affected 100% of reef-to-bay *Porites*, 50% of bay native *Porites,* and 20% of reef-to-bay *S. siderea* fragments during T12 only. Some *Porites* fragments were lost/missing from the bay site, likely due to the interference of shore fishing. During T4, 10% of reef-to-bay *Porites* fragments were missing and during T12, 50% of reef-to-bay *Porites* and 10% of bay native *Porites* were missing. These missing fragments were not included in partial mortality estimates.

**Supplementary Tables and Figures**

**Table S1**

Summary statistics for temperature (T, °C), photosynthetically active radiation (PAR, µmol m^-2^ s-^1^), pH_T_ (total scale), dissolved oxygen (DO, mg^-1^ L^-1^) concentrations, salinity, nitrate, ammonium, and phosphate concentrations (μmol L^-1^) at the transplant sites (Spanish Water Bay and Carmabi House Reef) and the source reef site (Tugboat Reef). SE = standard error, max = maximum, min = minimum, and N = number of days.

|  |  |  | Spanish Water Bay | | | Carmabi House Reef | | | Tugboat Reef | | |
| --- | --- | --- | --- | --- | --- | --- | --- | --- | --- | --- | --- |
| Site | | | Bay (transplant site) | | | Reef (transplant site) | | | Reef (source) | | |
| Timepoint | | | T0 | T4 | T12 | T0 | T4 | T12 | T0 | T4 | T12 |
| T°C | Daily Mean [SE] | | 28.8[0.04] | 29.47[0.04] | 29.59[0.07] | 27.66[0.04] | 29.38[0.03] | 28.3[0.05] | 27.93[0.02] | 29.41[0.02] | 28.29[0.05] |
|  | Mean Daily Range [SE] | | 1.02[0.03] | 0.76[0.05] | 1.03[0.05] | 0.6[0.04] | 0.47[0.03] | 0.63[0.02] | 0.61[0.02] | 0.69[0.03] | 0.69[0.02] |
|  | Max. Daily Range | | 1.30 | 1.40 | 1.70 | 1.10 | 0.80 | 0.90 | 0.80 | 0.90 | 1.00 |
|  | Max. |  | 29.90 | 30.50 | 31.04 | 28.40 | 30.00 | 29.10 | 28.50 | 30.10 | 29.20 |
|  | Min. |  | 27.71 | 28.70 | 28.10 | 26.50 | 28.90 | 27.00 | 27.40 | 28.80 | 26.90 |
|  | N |  | 30 | 29 | 31 | 30 | 32 | 35 | 33 | 31 | 46 |
| PAR | Daily Mean (SE) | | 441[14] | 259[24] | 513[16] | 354[12] | 303[19] | 434[15] | 343[11.25] | 288[35.12] | 444[8.16] |
|  | Max. |  | 1273 | 1013 | 1298 | 985 | 1056 | 1101 | 832 | 894 | 926 |
|  | N |  | 18 | 21 | 26 | 16 | 22 | 20 | 12 | 3 | 5 |
| pH_T_ | Daily Mean [SE] | | 8.01[0.01] | 7.95[0.01] | 8.05[0.01] | 8.00[0.01] | 8.03[0.01] | 8.08[0.02] | 8.00[0.01] | NA | 8.00[0.01] |
|  | Mean Daily Range [SE] | | 0.12[0.04] | 0.17[0.03] | 0.06[0.004] | 0.07[0.002] | 0.08[0.01] | 0.07[0.01] | 0.12[0.01] | NA | 0.07[0.01] |
|  | Max. Daily Range | | 0.33 | 0.34 | 0.08 | 0.07 | 0.13 | 0.16 | 0.15 | NA | 0.10 |
|  | Max. |  | 8.17 | 8.16 | 8.15 | 8.04 | 8.12 | 8.17 | 8.08 | NA | 8.10 |
|  | Min. |  | 7.83 | 7.82 | 7.93 | 7.91 | 7.97 | 7.91 | 7.83 | NA | 7.92 |
|  | N |  | 7 | 9 | 12 | 4 | 7 | 16 | 9 | NA | 10 |
| DO | Daily Mean [SE] | | 6.49[0.02] | 6[0.07] | 6.68[0.04] | 6.8[0.03] | 6.76[0.02] | 6.64[0.02] | 6.58[0.01] | 6.48[0.01] | 6.42[0.01] |
|  | Mean Daily Range [SE] | | 2[0.06] | 2.66[0.17] | 1.88[0.1] | 1.57[0.19] | 0.69[0.08] | 1.26[0.12] | 1.13[0.04] | 0.97[0.11] | 1.19[0.05] |
|  | Max. Daily Range | | 2.93 | 5.49 | 2.73 | 3.59 | 1.43 | 4.51 | 1.49 | 1.19 | 1.48 |
|  | Max. |  | 8.03 | 8.41 | 8.58 | 9.61 | 7.25 | 10.55 | 7.34 | 6.96 | 7.11 |
|  | Min. |  | 4.95 | 1.96 | 5.38 | 5.75 | 5.82 | 5.87 | 5.75 | 5.77 | 5.63 |
|  | N |  | 26 | 35 | 20 | 30 | 14 | 41 | 23 | 5 | 8 |
| Salinity | Daily Mean [SE] | | 36.11[0.27] | 34.71[0.11] | 36.41[0.21] | NA | 33.16[0.06] | 33.66[0.06] | 34.53[0.04] | 32.96[0.04] | 33.52[0.19] |
|  | Mean Daily Range [SE] | | 1.07[0.34] | 0.7[0.13] | 1.2[0.6] | NA | 0.49[0.13] | 1.92[0.15] | 0.59[0.1] | 0.3[0.06] | 2.54[0.54] |
|  | Max. Daily Range | | 6.50 | 3.89 | 9.54 | NA | 1.69 | 4.32 | 1.77 | 0.50 | 3.74 |
|  | Max. |  | 37.66 | 35.62 | 38.99 | NA | 33.77 | 35.02 | 35.02 | 33.37 | 36.44 |
|  | Min. |  | 29.70 | 30.76 | 29.24 | NA | 31.78 | 30.04 | 33.25 | 32.77 | 30.33 |
|  | N |  | 26 | 35 | 20 | NA | 14 | 52 | 23 | 5 | 8 |
| Nitrate | Mean[SE] | | 0.40[0.12] | 0.29[0.08] | 0.25[0.11] | 0.59[0.10] | 0.37[0.15] | 0.68[0.18] | NA | NA | 0.21[0.07] |
|  | N |  | 7 | 7 | 10 | 7 | 5 | 6 | NA | NA | 5 |
| Ammonium | Mean[SE] | | 2.65[0.54] | 2.11[0.31] | 1.98[0.51] | 2.75[0.73] | 1.40[0.47] | 2.63[0.78] | NA | NA | 1.46[0.59] |
|  | N |  | 8 | 7 | 10 | 6 | 4 | 6 | NA | NA | 5 |
| Phosphate | Mean[SE] | | 0.09[0.01] | 0.07[0.01] | 0.13[0.03] | 0.13[0.02] | 0.11[0.03] | 0.07[0.01] | NA | NA | 0.09[0.04] |
|  | N |  | 7 | 7 | 10 | 6 | 5 | 6 | NA | NA | 5 |

**Table S2**

Results from non-parametric tests (Scheirer-Ray-Hare extension of the Kruskal-Wallis and *post-hoc* Dunn’s tests) comparing daily average temperature, photosynthetic active radiation (PAR), pH_T_, dissolved oxygen (DO) concentrations, and salinity between all three sites (CH = Carmabi House Reef, TB = Tugboat Reef, SW = Spanish Water Bay) and timepoints (T0, T4, T12). df = degrees of freedom, SS =sum of squares, adj. p-value = p-value adjusted using Bonferroni correction. Only within timepoint significant p-values (<0.05) are bolded for pairwise comparisons.

| T °C (Daily Mean) |  |  |  |  |  |  |  |
| --- | --- | --- | --- | --- | --- | --- | --- |
| Effect | **df** | **SS** | **H-value** | **p-value** | **Effect for pairwise comparisons** | **Pairwise comparisons** | **adj. p-value** |
| Site | 2 | 553452 | 75.04 | **0** | Site x Timepoint | CH_T0 - CH_T12 | **1.90 × 10⁻²** |
| Timepoint | 2 | 1140024 | 154.57 | **0** |  | CH_T0 - CH_T4 | **2.25 × 10⁻¹⁸** |
| Site x Timepoint | 4 | 208668 | 28.29 | **1.09 x 10^-5^** |  | CH_T12 - CH_T4 | **8.49 × 10⁻⁸** |
|  |  |  |  |  |  | CH_T0 - TB_T0 | 1 |
|  |  |  |  |  |  | CH_T12 - TB_T0 | 6.33 × 10⁻¹ |
|  |  |  |  |  |  | CH_T4 - TB_T0 | **8.07 × 10⁻¹⁵** |
|  |  |  |  |  |  | CH_T0 - TB_T12 | **5.37 × 10⁻³** |
|  |  |  |  |  |  | CH_T12 - TB_T12 | 1 |
|  |  |  |  |  |  | CH_T4 - TB_T12 | **1.74 × 10⁻⁸** |
|  |  |  |  |  |  | TB_T0 - TB_T12 | 2.93 × 10⁻¹ |
|  |  |  |  |  |  | CH_T0 - TB_T4 | **2.55 × 10⁻¹⁹** |
|  |  |  |  |  |  | CH_T12 - TB_T4 | **1.62 × 10⁻⁸** |
|  |  |  |  |  |  | CH_T4 - TB_T4 | 1 |
|  |  |  |  |  |  | TB_T0 - TB_T4 | **1.03 × 10⁻¹⁵** |
|  |  |  |  |  |  | TB_T12 - TB_T4 | **2.90 × 10⁻⁹** |
|  |  |  |  |  |  | CH_T0 - SW_T0 | **1.49 × 10⁻⁷** |
|  |  |  |  |  |  | CH_T12 - SW_T0 | 3.03 × 10⁻¹ |
|  |  |  |  |  |  | CH_T4 - SW_T0 | 5.54 × 10⁻² |
|  |  |  |  |  |  | TB_T0 - SW_T0 | **3.78 × 10⁻⁵** |
|  |  |  |  |  |  | TB_T12 - SW_T0 | 2.69 × 10⁻¹ |
|  |  |  |  |  |  | TB_T4 - SW_T0 | **2.05 × 10⁻²** |
|  |  |  |  |  |  | CH_T0 - SW_T12 | **5.44 × 10⁻²²** |
|  |  |  |  |  |  | CH_T12 - SW_T12 | **2.03 × 10⁻¹⁰** |
|  |  |  |  |  |  | CH_T4 - SW_T12 | 1 |
|  |  |  |  |  |  | TB_T0 - SW_T12 | **3.38 × 10⁻¹⁸** |
|  |  |  |  |  |  | TB_T12 - SW_T12 | **2.29 × 10⁻¹¹** |
|  |  |  |  |  |  | TB_T4 - SW_T12 | 1 |
|  |  |  |  |  |  | SW_T0 - SW_T12 | **1.66 × 10⁻³** |
|  |  |  |  |  |  | CH_T0 - SW_T4 | **2.82 ×10⁻²⁰** |
|  |  |  |  |  |  | CH_T12 - SW_T4 | **2.47 × 10⁻⁹** |
|  |  |  |  |  |  | CH_T4 - SW_T4 | 1 |
|  |  |  |  |  |  | TB_T0 - SW_T4 | **1.20 × 10⁻¹⁶** |
|  |  |  |  |  |  | TB_T12 - SW_T4 | **3.97 × 10⁻¹⁰** |
|  |  |  |  |  |  | TB_T4 - SW_T4 | 1 |
|  |  |  |  |  |  | SW_T0 - SW_T4 | **5.79 × 10⁻³** |
|  |  |  |  |  |  | SW_T12 - SW_T4 | 1 |
| T °C (Mean Daily Range) |  |  |  |  |  |  |  |
| Effect | **df** | **SS** | **H-value** | **p-value** | **Effect for pairwise comparisons** | **Pairwise comparisons** | **adj. p-value** |
| Site | 2 | 762452 | 104.135 | 0 | Site x Timepoint | CH_T0 - CH_T12 | 1 |
| Timepoint | 2 | 77195 | 10.543 | **5.14 x 10^-3^** |  | CH_T0 - CH_T4 | 1 |
| Site x Timepoint | 4 | 133529 | 18.237 | **1.11 x 10^-3^** |  | CH_T12 - CH_T4 | 7.72 × 10⁻¹ |
|  |  |  |  |  |  | CH_T0 - TB_T0 | 1 |
|  |  |  |  |  |  | CH_T12 - TB_T0 | 1 |
|  |  |  |  |  |  | CH_T4 - TB_T0 | 1 |
|  |  |  |  |  |  | CH_T0 - TB_T12 | 1 |
|  |  |  |  |  |  | CH_T12 - TB_T12 | 1 |
|  |  |  |  |  |  | CH_T4 - TB_T12 | **1.81 × 10⁻³** |
|  |  |  |  |  |  | TB_T0 - TB_T12 | 1 |
|  |  |  |  |  |  | CH_T0 - TB_T4 | 1 |
|  |  |  |  |  |  | CH_T12 - TB_T4 | 1 |
|  |  |  |  |  |  | CH_T4 - TB_T4 | **4.76 × 10⁻³** |
|  |  |  |  |  |  | TB_T0 - TB_T4 | 1 |
|  |  |  |  |  |  | TB_T12 - TB_T4 | 1 |
|  |  |  |  |  |  | CH_T0 - SW_T0 | **2.02 × 10⁻¹⁰** |
|  |  |  |  |  |  | CH_T12 - SW_T0 | **1.28 × 10⁻⁹** |
|  |  |  |  |  |  | CH_T4 - SW_T0 | **1.23 × 10⁻¹⁶** |
|  |  |  |  |  |  | TB_T0 - SW_T0 | **4.52 × 10⁻¹⁰** |
|  |  |  |  |  |  | TB_T12 - SW_T0 | **1.91 × 10⁻⁶** |
|  |  |  |  |  |  | TB_T4 - SW_T0 | **4.05 × 10⁻⁵** |
|  |  |  |  |  |  | CH_T0 - SW_T12 | **2.75 × 10⁻⁸** |
|  |  |  |  |  |  | CH_T12 - SW_T12 | **1.70 × 10⁻⁷** |
|  |  |  |  |  |  | CH_T4 - SW_T12 | **6.00 × 10⁻¹⁴** |
|  |  |  |  |  |  | TB_T0 - SW_T12 | **6.30 × 10⁻⁸** |
|  |  |  |  |  |  | TB_T12 - SW_T12 | **1.37 × 10⁻⁴** |
|  |  |  |  |  |  | TB_T4 - SW_T12 | **1.43 × 10⁻³** |
|  |  |  |  |  |  | SW_T0 - SW_T12 | 1 |
|  |  |  |  |  |  | CH_T0 - SW_T4 | 2.02 × 10⁻¹ |
|  |  |  |  |  |  | CH_T12 - SW_T4 | 6.77 × 10⁻¹ |
|  |  |  |  |  |  | CH_T4 - SW_T4 | **2.50 × 10⁻⁴** |
|  |  |  |  |  |  | TB_T0 - SW_T4 | 3.79 × 10⁻¹ |
|  |  |  |  |  |  | TB_T12 - SW_T4 | 1 |
|  |  |  |  |  |  | TB_T4 - SW_T4 | 1 |
|  |  |  |  |  |  | SW_T0 - SW_T4 | **1.76 × 10⁻³** |
|  |  |  |  |  |  | SW_T12 - SW_T4 | **3.39 × 10⁻²** |
| PAR (Daily Mean) |  |  |  |  |  |  |  |
| Effect | **df** | **SS** | **H-value** | **p-value** | **Effect for pairwise comparisons** | **Pairwise comparisons** | **adj. p-value** |
| Site | 2 | 13684 | 7.974 | **1.90 x 10^-2^** | Site | CH - TB | 1 |
| Timepoint | 2 | 122836 | 71.583 | 0 |  | CH - SW | **1.97 × 10⁻²** |
| Site x Timepoint | 4 | 15813 | 9.215 | 5.59 x 10^-2^ |  | TB - SW | 5.73 × 10⁻² |
| pH_T_ (Daily Mean) |  |  |  |  |  |  |  |
| Effect | **df** | **SS** | **H-value** | **p-value** | **Effect for pairwise comparisons** | **Pairwise comparisons** | **adj. p-value** |
| Site | 2 | 5609 | 12.1276 | **2.33 × 10⁻³** | Site x Timepoint | CH_T0 - CH_T12 | 1.89 × 10⁻¹ |
| Timepoint | 2 | 8651.1 | 18.705 | **8.67 × 10⁻⁵** |  | CH_T0 - CH_T4 | 1 |
| Site x Timepoint | 3 | 4473.9 | 9.6732 | **2.16 × 10⁻²** |  | CH_T12 - CH_T4 | 1 |
|  |  |  |  |  |  | CH_T0 - TB_T0 | 1 |
|  |  |  |  |  |  | CH_T12 - TB_T0 | **2.93 × 10⁻²** |
|  |  |  |  |  |  | CH_T4 - TB_T0 | 1 |
|  |  |  |  |  |  | CH_T0 - TB_T12 | 1 |
|  |  |  |  |  |  | CH_T12 - TB_T12 | **3.45 × 10⁻²** |
|  |  |  |  |  |  | CH_T4 - TB_T12 | 1 |
|  |  |  |  |  |  | TB_T0 - TB_T12 | 1 |
|  |  |  |  |  |  | CH_T0 - SW_T0 | 1 |
|  |  |  |  |  |  | CH_T12 - SW_T0 | 3.75 × 10⁻¹ |
|  |  |  |  |  |  | CH_T4 - SW_T0 | 1 |
|  |  |  |  |  |  | TB_T0 - SW_T0 | 1 |
|  |  |  |  |  |  | TB_T12 - SW_T0 | 1 |
|  |  |  |  |  |  | CH_T0 - SW_T12 | 1 |
|  |  |  |  |  |  | CH_T12 - SW_T12 | 1 |
|  |  |  |  |  |  | CH_T4 - SW_T12 | 1 |
|  |  |  |  |  |  | TB_T0 - SW_T12 | 7.06 × 10⁻¹ |
|  |  |  |  |  |  | TB_T12 - SW_T12 | 8.67 × 10⁻¹ |
|  |  |  |  |  |  | SW_T0 - SW_T12 | 1 |
|  |  |  |  |  |  | CH_T0 - SW_T4 | 1 |
|  |  |  |  |  |  | CH_T12 - SW_T4 | **4.28 × 10⁻⁷** |
|  |  |  |  |  |  | CH_T4 - SW_T4 | **2.85 × 10⁻²** |
|  |  |  |  |  |  | TB_T0 - SW_T4 | 9.90 × 10⁻¹ |
|  |  |  |  |  |  | TB_T12 - SW_T4 | 6.07 × 10⁻¹ |
|  |  |  |  |  |  | SW_T0 - SW_T4 | 3.95 × 10⁻¹ |
|  |  |  |  |  |  | SW_T12 - SW_T4 | **2.02 × 10⁻⁴** |
| pH_T_ (Mean Daily Range) |  |  |  |  |  |  |  |
| Effect | **df** | **SS** | **H-value** | **p-value** | **Effect for pairwise comparisons** | **Pairwise comparisons** | **adj. p-value** |
| Site | 2 | 1887.7 | 4.0815 | 1.30 × 10⁻¹ | Timepoint | T0 - T12 | **6.04 × 10⁻³** |
| Timepoint | 2 | 7775.8 | 16.8125 | **2.23 × 10⁻⁴** |  | T0 - T4 | 1 |
| Site x Timepoint | 3 | 3211.7 | 6.9441 | 7.37 × 10⁻² |  | T12 - T4 | **9.55 × 10⁻⁴** |
| DO (Daily Mean) |  |  |  |  |  |  |  |
| Effect | **df** | **SS** | **H-value** | **p-value** | **Effect for pairwise comparisons** | **Pairwise comparisons** | **adj. p-value** |
| Site | 2 | 201123 | 58.857 | **0** | Site x Timepoint | CH_T0 - CH_T12 | 7.00 × 10⁻² |
| Timepoint | 2 | 50792 | 14.864 | **5.92 x 10^-4^** |  | CH_T0 - CH_T4 | 1 |
| Site x Timepoint | 4 | 160370 | 46.931 | **0** |  | CH_T12 - CH_T4 | 3.79 × 10⁻¹ |
|  |  |  |  |  |  | CH_T0 - TB_T0 | **4.13 × 10⁻³** |
|  |  |  |  |  |  | CH_T12 - TB_T0 | 1 |
|  |  |  |  |  |  | CH_T4 - TB_T0 | **3.57 × 10⁻²** |
|  |  |  |  |  |  | CH_T0 - TB_T12 | **9.93 × 10⁻⁶** |
|  |  |  |  |  |  | CH_T12 - TB_T12 | **2.76 × 10⁻²** |
|  |  |  |  |  |  | CH_T4 - TB_T12 | **8.46 × 10⁻⁵** |
|  |  |  |  |  |  | TB_T0 - TB_T12 | 6.27 × 10⁻¹ |
|  |  |  |  |  |  | CH_T0 - TB_T4 | **1.18 × 10⁻²** |
|  |  |  |  |  |  | CH_T12 - TB_T4 | 1 |
|  |  |  |  |  |  | CH_T4 - TB_T4 | **2.23 × 10⁻²** |
|  |  |  |  |  |  | TB_T0 - TB_T4 | 1 |
|  |  |  |  |  |  | TB_T12 - TB_T4 | 1 |
|  |  |  |  |  |  | CH_T0 - SW_T0 | **4.70 × 10⁻⁷** |
|  |  |  |  |  |  | CH_T12 - SW_T0 | 6.81 × 10⁻² |
|  |  |  |  |  |  | CH_T4 - SW_T0 | **7.77 × 10⁻⁵** |
|  |  |  |  |  |  | TB_T0 - SW_T0 | 1 |
|  |  |  |  |  |  | TB_T12 - SW_T0 | 1 |
|  |  |  |  |  |  | TB_T4 - SW_T0 | 1 |
|  |  |  |  |  |  | CH_T0 - SW_T12 | 1 |
|  |  |  |  |  |  | CH_T12 - SW_T12 | 1 |
|  |  |  |  |  |  | CH_T4 - SW_T12 | 1 |
|  |  |  |  |  |  | TB_T0 - SW_T12 | 1 |
|  |  |  |  |  |  | TB_T12 - SW_T12 | **8.90 × 10⁻³** |
|  |  |  |  |  |  | TB_T4 - SW_T12 | 5.17 × 10⁻¹ |
|  |  |  |  |  |  | SW_T0 - SW_T12 | **2.42 × 10⁻²** |
|  |  |  |  |  |  | CH_T0 - SW_T4 | **1.62 × 10⁻¹⁹** |
|  |  |  |  |  |  | CH_T12 - SW_T4 | **1.32 × 10⁻¹⁰** |
|  |  |  |  |  |  | CH_T4 - SW_T4 | **1.44 × 10⁻¹²** |
|  |  |  |  |  |  | TB_T0 - SW_T4 | **7.36 × 10⁻⁵** |
|  |  |  |  |  |  | TB_T12 - SW_T4 | 1 |
|  |  |  |  |  |  | TB_T4 - SW_T4 | 1 |
|  |  |  |  |  |  | SW_T0 - SW_T4 | 5.51 × 10⁻² |
|  |  |  |  |  |  | SW_T12 - SW_T4 | **2.28 × 10⁻⁹** |
| DO (Mean Daily Range) |  |  |  |  |  |  |  |
| Effect | **df** | **SS** | **H-value** | **p-value** | **Effect for pairwise comparisons** | **Pairwise comparisons** | **adj. p-value** |
| Site | 2 | 289362 | 84.679 | **0** | Site x Timepoint | CH_T0 - CH_T12 | 1 |
| Timepoint | 2 | 5522 | 1.616 | 4.46 x 10^-1^ |  | CH_T0 - CH_T4 | **2.35 × 10⁻²** |
| Site x Timepoint | 4 | 45961 | 13.45 | **9.27 x 10^-3^** |  | CH_T12 - CH_T4 | 3.99 × 10⁻¹ |
|  |  |  |  |  |  | CH_T0 - TB_T0 | 1 |
|  |  |  |  |  |  | CH_T12 - TB_T0 | 1 |
|  |  |  |  |  |  | CH_T4 - TB_T0 | 8.40 × 10⁻¹ |
|  |  |  |  |  |  | CH_T0 - TB_T12 | 1 |
|  |  |  |  |  |  | CH_T12 - TB_T12 | 1 |
|  |  |  |  |  |  | CH_T4 - TB_T12 | 1 |
|  |  |  |  |  |  | TB_T0 - TB_T12 | 1 |
|  |  |  |  |  |  | CH_T0 - TB_T4 | 1 |
|  |  |  |  |  |  | CH_T12 - TB_T4 | 1 |
|  |  |  |  |  |  | CH_T4 - TB_T4 | 1 |
|  |  |  |  |  |  | TB_T0 - TB_T4 | 1 |
|  |  |  |  |  |  | TB_T12 - TB_T4 | 1 |
|  |  |  |  |  |  | CH_T0 - SW_T0 | **1.83 × 10⁻²** |
|  |  |  |  |  |  | CH_T12 - SW_T0 | **2.29 × 10⁻⁵** |
|  |  |  |  |  |  | CH_T4 - SW_T0 | **3.02 × 10⁻⁸** |
|  |  |  |  |  |  | TB_T0 - SW_T0 | **3.52 × 10⁻⁴** |
|  |  |  |  |  |  | TB_T12 - SW_T0 | 1.83 × 10⁻¹ |
|  |  |  |  |  |  | TB_T4 - SW_T0 | **4.20 × 10⁻²** |
|  |  |  |  |  |  | CH_T0 - SW_T12 | 2.82 × 10⁻¹ |
|  |  |  |  |  |  | CH_T12 - SW_T12 | **2.52 × 10⁻³** |
|  |  |  |  |  |  | CH_T4 - SW_T12 | **2.86 × 10⁻⁶** |
|  |  |  |  |  |  | TB_T0 - SW_T12 | **1.13 × 10⁻²** |
|  |  |  |  |  |  | TB_T12 - SW_T12 | 7.41 × 10⁻¹ |
|  |  |  |  |  |  | TB_T4 - SW_T12 | 1.61 × 10⁻¹ |
|  |  |  |  |  |  | SW_T0 - SW_T12 | 1 |
|  |  |  |  |  |  | CH_T0 - SW_T4 | **3.91 × 10⁻⁵** |
|  |  |  |  |  |  | CH_T12 - SW_T4 | **1.06 × 10⁻⁹** |
|  |  |  |  |  |  | CH_T4 - SW_T4 | **8.62 × 10⁻¹²** |
|  |  |  |  |  |  | TB_T0 - SW_T4 | **2.94 × 10⁻⁷** |
|  |  |  |  |  |  | TB_T12 - SW_T4 | **1.11 × 10⁻²** |
|  |  |  |  |  |  | TB_T4 - SW_T4 | **3.39 × 10⁻³** |
|  |  |  |  |  |  | SW_T0 - SW_T4 | 1 |
|  |  |  |  |  |  | SW_T12 - SW_T4 | 1 |
| Salinity (Daily Mean) |  |  |  |  |  |  |  |
| Effect | **df** | **SS** | **H-value** | **p-value** | **Effect for pairwise comparisons** | **Pairwise comparisons** | **adj. p-value** |
| Site | 2 | 245508 | 87.494 | **0** | Site x Timepoint | CH_T12 - CH_T4 | 1 |
| Timepoint | 2 | 61541 | 21.932 | **1.70 x 10^-5^** |  | CH_T12 - TB_T0 | **1.25 × 10⁻³** |
| Site x Timepoint | 3 | 23841 | 8.496 | **3.68 x 10^-2^** |  | CH_T4 - TB_T0 | **3.86 × 10⁻⁵** |
|  |  |  |  |  |  | CH_T12 - TB_T12 | 1 |
|  |  |  |  |  |  | CH_T4 - TB_T12 | 1 |
|  |  |  |  |  |  | TB_T0 - TB_T12 | 1.39 × 10⁻¹ |
|  |  |  |  |  |  | CH_T12 - TB_T4 | 1 |
|  |  |  |  |  |  | CH_T4 - TB_T4 | 1 |
|  |  |  |  |  |  | TB_T0 - TB_T4 | **7.71 × 10⁻³** |
|  |  |  |  |  |  | TB_T12 - TB_T4 | 1 |
|  |  |  |  |  |  | CH_T12 - SW_T0 | **2.70 × 10⁻¹¹** |
|  |  |  |  |  |  | CH_T4 - SW_T0 | **6.07 × 10⁻¹¹** |
|  |  |  |  |  |  | TB_T0 - SW_T0 | 4.39 × 10⁻¹ |
|  |  |  |  |  |  | TB_T12 - SW_T0 | **1.41 × 10⁻⁴** |
|  |  |  |  |  |  | TB_T4 - SW_T0 | **9.92 × 10⁻⁶** |
|  |  |  |  |  |  | CH_T12 - SW_T12 | **8.53 × 10⁻¹¹** |
|  |  |  |  |  |  | CH_T4 - SW_T12 | **5.79 × 10⁻¹¹** |
|  |  |  |  |  |  | TB_T0 - SW_T12 | 2.19 × 10⁻¹ |
|  |  |  |  |  |  | TB_T12 - SW_T12 | **7.24 × 10⁻⁵** |
|  |  |  |  |  |  | TB_T4 - SW_T12 | **5.12 × 10⁻⁶** |
|  |  |  |  |  |  | SW_T0 - SW_T12 | 1 |
|  |  |  |  |  |  | CH_T12 - SW_T4 | **1.05 × 10⁻⁵** |
|  |  |  |  |  |  | CH_T4 - SW_T4 | **1.37 × 10⁻⁶** |
|  |  |  |  |  |  | TB_T0 - SW_T4 | 1 |
|  |  |  |  |  |  | TB_T12 - SW_T4 | 4.27 × 10⁻² |
|  |  |  |  |  |  | TB_T4 - SW_T4 | **2.29 × 10⁻³** |
|  |  |  |  |  |  | SW_T0 - SW_T4 | 5.56 × 10⁻¹ |
|  |  |  |  |  |  | SW_T12 - SW_T4 | 2.73 × 10⁻¹ |
| Salinity (Mean Daily Range) |  |  |  |  |  |  |  |
| Effect | **df** | **SS** | **H-value** | **p-value** | **Effect for pairwise comparisons** | **Pairwise comparisons** | **adj. p-value** |
| Site | 2 | 10061 | 3.5858 | 1.66 × 10⁻¹ | Site x Timepoint | CH_T12 - CH_T4 | **1.27 × 10⁻⁵** |
| Timepoint | 2 | 53401 | 19.0317 | **7.40 × 10⁻⁵** |  | CH_T12 - TB_T0 | **3.86 × 10⁻⁴** |
| Site x Timepoint | 3 | 45089 | 16.0694 | **1.10 × 10⁻³** |  | CH_T4 - TB_T0 | 1 |
|  |  |  |  |  |  | CH_T12 - TB_T12 | 1 |
|  |  |  |  |  |  | CH_T4 - TB_T12 | **2.04 × 10⁻²** |
|  |  |  |  |  |  | TB_T0 - TB_T12 | 2.61 × 10⁻¹ |
|  |  |  |  |  |  | CH_T12 - TB_T4 | **1.41 × 10⁻²** |
|  |  |  |  |  |  | CH_T4 - TB_T4 | 1 |
|  |  |  |  |  |  | TB_T0 - TB_T4 | 1 |
|  |  |  |  |  |  | TB_T12 - TB_T4 | 1.35 × 10⁻¹ |
|  |  |  |  |  |  | CH_T12 - SW_T0 | **1.38 × 10⁻⁴** |
|  |  |  |  |  |  | CH_T4 - SW_T0 | 1 |
|  |  |  |  |  |  | TB_T0 - SW_T0 | 1 |
|  |  |  |  |  |  | TB_T12 - SW_T0 | 2.18 × 10⁻¹ |
|  |  |  |  |  |  | TB_T4 - SW_T0 | 1 |
|  |  |  |  |  |  | CH_T12 - SW_T12 | **1.68 × 10⁻³** |
|  |  |  |  |  |  | CH_T4 - SW_T12 | 1 |
|  |  |  |  |  |  | TB_T0 - SW_T12 | 1 |
|  |  |  |  |  |  | TB_T12 - SW_T12 | 3.75 × 10⁻¹ |
|  |  |  |  |  |  | TB_T4 - SW_T12 | 1 |
|  |  |  |  |  |  | SW_T0 - SW_T12 | 1 |
|  |  |  |  |  |  | CH_T12 - SW_T4 | **3.77 × 10⁻⁵** |
|  |  |  |  |  |  | CH_T4 - SW_T4 | 1 |
|  |  |  |  |  |  | TB_T0 - SW_T4 | 1 |
|  |  |  |  |  |  | TB_T12 - SW_T4 | 2.31 × 10⁻¹ |
|  |  |  |  |  |  | TB_T4 - SW_T4 | 1 |
|  |  |  |  |  |  | SW_T0 - SW_T4 | 1 |
|  |  |  |  |  |  | SW_T12 - SW_T4 | 1 |
| Nitrate |  |  |  |  |  |  |  |
| Effect | **df** | **SS** | **H-value** | **p-value** | **Effect for pairwise comparisons** | **Pairwise comparisons** | **adj. p-value** |
| Site | 2 | 1169.8 | 6.2246 | **4.45 × 10⁻²** | Site | CH - TB | 1.43 × 10⁻¹ |
| Timepoint | 2 | 371.1 | 1.975 | 3.73 × 10⁻¹ |  | CH - SW | **3.72 × 10⁻²** |
| Site x Timepoint | 2 | 463.7 | 2.4677 | 2.91 × 10⁻¹ |  | TB - SW | 1 |
| Ammonium |  |  |  |  |  |  |  |
| Effect | **df** | **SS** | **H-value** | **p-value** | **Effect for pairwise comparisons** | **Pairwise comparisons** | **adj. p-value** |
| Site | 2 | 252.2 | 1.4001 | 4.97 × 10⁻¹ |  |  |  |
| Timepoint | 2 | 407.2 | 2.2605 | 3.23 × 10⁻¹ |  |  |  |
| Site x Timepoint | 2 | 295.2 | 1.6387 | 4.41 × 10⁻¹ |  |  |  |
| Phosphate |  |  |  |  |  |  |  |
| Effect | **df** | **SS** | **H-value** | **p-value** | **Effect for pairwise comparisons** | **Pairwise comparisons** | **adj. p-value** |
| Site | 2 | 271.5 | 1.5078 | 4.71 × 10⁻¹ |  |  |  |
| Timepoint | 2 | 500.2 | 2.7779 | 2.49 × 10⁻¹ |  |  |  |
| Site x Timepoint | 2 | 386.7 | 2.1476 | 3.42 × 10⁻¹ |  |  |  |

**Table S3**

Temperature (°C) and photosynthetically active radiation (PAR, µmol m^-1^ s^-1^) of photosynthesis to respiration incubations after 0, 4, and 12 months of transplantation.

| Timepoint | Transplant site | Temperature | PAR |
| --- | --- | --- | --- |
| T0 | Bay | 28.5 | 310 |
|  | Reef | 28.0 | 520 |
| T4 | Bay | 30.0 | 275 |
|  | Reef | 29.5 | 450 |
| T12 | Bay | 28.5 | 350 |
|  | Reef | 28.0 | 520 |

**Table S4**

Results of (G)LMMs to assess effects of origin (bay or reef), treatment (native or transplant), and timepoint (T0, T4, T12) on daily gross photosynthesis: daily respiration (P_G day_/R_day_), chlorophyll a concentrations, symbiont density, tissue biomass, calcification rates, gross photosynthesis (P_G_), dark respiration (R_dark_), and light-enhanced dark respiration (LEDR). (G)LMM = (Generalized) Linear mixed model, SS = Sum of squares, MS = Mean squares, Num df = numerator degrees of freedom, Den df = denominator degrees of freedom, Chisq = Chi-squared statistic, npar = number of parameters, logLik = log-liklihood, AIC = Akaike Information Criterion, LRT = likelihood ratio test, Df = degrees of freedom. For *post-hoc* results, see Table S5.

| ***Siderastrea siderea*** | | | | | | | |
| --- | --- | --- | --- | --- | --- | --- | --- |
| **P_G day_/R_day_** | |  |  |  |  |  |  |
|  | LMM | SS | MS | Num df | Den df | F statistic | p-value |
|  | timepoint | 0.151 | 0.0753 | 2 | 87.590 | 2.297 | 0.107 |
|  | treatment | 0.0566 | 0.0566 | 1 | 87.425 | 1.726 | 0.192 |
|  | origin | 0.00783 | 0.00783 | 1 | 18.164 | 0.239 | 0.631 |
|  | timepoint x treatment | 0.0398 | 0.0199 | 2 | 88.042 | 0.606 | 0.548 |
|  | timepoint x origin | 1.687 | 0.844 | 2 | 87.590 | 25.725 | **<0.0001** |
|  | treatment x origin | 0.419 | 0.419 | 1 | 87.425 | 12.764 | **0.000578** |
|  | timepoint x treatment x origin | 0.220 | 0.110 | 2 | 88.042 | 3.356 | **0.0394** |
|  | Random effects | npar | logLik | AIC | LRT | Df | p-value |
|  | reduced | 14 | 10.733 | 6.5 |  |  |  |
|  | full | 13 | 8.745 | 8.5 | 3.975 | 1 | **0.0462** |
| **Chlorophyll a (µg cm^-2^)** | |  |  |  |  |  |  |
|  | LMM | SS | MS | Num df | Den df | F statistic | p-value |
|  | timepoint | 1.229 | 0.615 | 2 | 70.717 | 67.068 | **<0.0001** |
|  | treatment | 0.0955 | 0.00955 | 1 | 71.405 | 1.042 | 0.311 |
|  | origin | 0.0199 | 0.0199 | 1 | 18.113 | 2.177 | 0.157 |
|  | timepoint x treatment | 0.00580 | 0.0290 | 2 | 70.339 | 0.317 | 0.730 |
|  | timepoint x origin | 0.218 | 0.109 | 2 | 70.717 | 11.895 | **<0.0001** |
|  | treatment x origin | 0.0269 | 0.0269 | 1 | 71.405 | 2.939 | 0.0908 |
|  | timepoint x treatment x origin | 0.0403 | 0.0201 | 2 | 70.339 | 2.197 | 0.119 |
|  | Random effects | npar | logLik | AIC | LRT | Df | p-value |
|  | reduced | 14 | 60.0 | -92.1 |  |  |  |
|  | full | 13 | 57.2 | -88.5 | 5.587 | 1 | **0.0181** |
| **Symbiont density (x10^6^ cells cm^-2^)** | |  |  |  |  |  |  |
|  | LMM | SS | MS | Num df | Den df | F statistic | p-value |
|  | timepoint | 25.217 | 12.609 | 2 | 68.134 | 52.501 | **<0.0001** |
|  | treatment | 0.416 | 0.416 | 1 | 68.792 | 1.732 | 0.193 |
|  | origin | 0.304 | 0.304 | 1 | 17.172 | 1.265 | 0.276 |
|  | timepoint x treatment | 0.257 | 0.129 | 2 | 67.650 | 0.535 | 0.588 |
|  | timepoint x origin | 0.267 | 0.133 | 2 | 68.134 | 0.555 | 0.577 |
|  | treatment x origin | 0.681 | 0.681 | 1 | 68.792 | 2.836 | 0.0967 |
|  | timepoint x treatment x origin | 0.481 | 0.241 | 2 | 67.650 | 1.001 | 0.373 |
|  | Random effects | npar | logLik | AIC | LRT | Df | p-value |
|  | reduced | 14 | -80.8 | 197.6 |  |  |  |
|  | full | 13 | -92.5 | 210.9 | 15.3 | 1 | **<0.0001** |
| **Tissue biomass (mg cm^-2^)** | |  |  |  |  |  |  |
|  | LMM | SS | MS | Num df | Den df | F statistic | p-value |
|  | timepoint | 4782.900 | 2391.400 | 2 | 72.519 | 6.685 | **0.002** |
|  | treatment | 141.200 | 141.200 | 1 | 73.584 | 0.395 | 0.532 |
|  | origin | 3791.400 | 3791.400 | 1 | 19.355 | 10.599 | **0.004** |
|  | timepoint x treatment | 1285.900 | 642.950 | 2 | 73.137 | 1.797 | 0.173 |
|  | timepoint x origin | 121.300 | 60.650 | 2 | 72.519 | 0.170 | 0.844 |
|  | treatment x origin | 206.300 | 206.300 | 1 | 73.584 | 0.577 | 0.450 |
|  | timepoint x treatment x origin | 2440.400 | 1220.200 | 2 | 73.137 | 3.411 | **0.0383** |
|  | Random effects | npar | logLik | AIC | LRT | Df | p-value |
|  | reduced | 14 | -392.0 | 811.99 |  |  |  |
|  | full | 13 | -393.0 | 812.5 | 2.51 | 1 | 0.113 |
| **Calcification (mg cm^-2^ d^-1^)** | |  |  |  |  |  |  |
|  | LMM | SS | MS | Num df | Den df | F statistic | p-value |
|  | timepoint | 0.726 | 0.726 | 1 | 51.856 | 7.642 | 0.00788 |
|  | treatment | 0.276 | 0.276 | 1 | 51.856 | 2.905 | 0.0943 |
|  | origin | 0.515 | 0.515 | 1 | 17.720 | 5.426 | **0.0319** |
|  | timepoint x treatment | 0.127 | 0.127 | 1 | 52.932 | 1.333 | 0.253 |
|  | timepoint x origin | 0.295 | 0.295 | 1 | 51.856 | 3.103 | 0.0840 |
|  | treatment x origin | 1.301 | 1.301 | 1 | 51.856 | 13.697 | **0.000521** |
|  | timepoint x treatment x origin | 0.206 | 0.206 | 1 | 52.932 | 2.170 | 0.147 |
|  | Random effects | npar | logLik | AIC | LRT | Df | p-value |
|  | reduced | 10 | -35.5 | 90.9 |  |  |  |
|  | full | 9 | -39.6 | 97.3 | 8.34 | 1 | **0.00387** |
| **P_GROSS_ (μmol O2 cm^-2^ h^-1^)** | |  |  |  |  |  |  |
|  | LMM | SS | MS | Num df | Den df | F statistic | p-value |
|  | timepoint | 0.508 | 0.254 | 2 | 87.618 | 8.0265 | **0.000630** |
|  | treatment | 0.0679 | 0.0679 | 1 | 87.463 | 2.144 | 0.147 |
|  | origin | 0.182 | 0.182 | 1 | 18.227 | 5.759 | **0.0273** |
|  | timepoint x treatment | 0.0710 | 0.0355 | 2 | 88.046 | 1.120 | 0.331 |
|  | timepoint x origin | 1.400 | 0.700 | 2 | 87.618 | 22.0994 | **<0.0001** |
|  | treatment x origin | 0.140 | 0.140 | 1 | 87.463 | 4.423 | **0.0383** |
|  | timepoint x treatment x origin | 0.188 | 0.0938 | 2 | 88.046 | 2.963 | 0.0568 |
|  | Random effects | npar | logLik | AIC | LRT | Df | p-value |
|  | reduced | 14 | 11.781 | 4.4 |  |  |  |
|  | full | 13 | 9.197 | 7.6 | 5.167 | 1 | 0.0230 |
| **R_DARK_ (μmol O2 cm^-2^ h^-1^)** | |  |  |  |  |  |  |
|  | GLMM | SS | MS | Chisq | Df | F statistic | p-value |
|  | timepoint | 1.461 | 0.731 | 21.863 | 2 | 12.828 | **<0.0001** |
|  | treatment | 0.0288 | 0.0288 | 0 | 1 | 0.505 | 1.000 |
|  | origin | 0.374 | 0.374 | 0.426 | 1 | 6.571 | 0.514 |
|  | timepoint x treatment | 0.199 | 0.0994 | 3.711 | 2 | 1.746 | 0.156 |
|  | timepoint x origin | 0.191 | 0.0956 | 3.350 | 2 | 1.679 | 0.187 |
|  | treatment x origin | 0.00246 | 0.00246 | 0 | 1 | 0.0433 | 1.000 |
|  | timepoint x treatment x origin | 0.0217 | 0.0108 | 0.452 | 2 | 0.190 | 0.798 |
|  | Random effects | npar | logLik | AIC | Chisq | Df | p-value |
|  | reduced | 13 | 28.3 | -30.6 |  |  |  |
|  | full | 14 | 34.6 | -41.2 | 12.6 | 1 | **0.000383** |
| **LEDR (μmol O2 cm^-2^ h^-1^)** | |  |  |  |  |  |  |
|  | LMM | SS | MS | Num df | Den df | F statistic | p-value |
|  | timepoint | 0.311 | 0.155 | 2 | 87.680 | 11.479 | <0.0001 |
|  | treatment | 0.00967 | 0.00967 | 1 | 87.482 | 0.714 | 0.400 |
|  | origin | 0.108 | 0.108 | 1 | 18.112 | 7.973 | **0.0112** |
|  | timepoint x treatment | 0.0213 | 0.0107 | 2 | 88.212 | 0.789 | 0.458 |
|  | timepoint x origin | 0.212 | 0.106 | 2 | 87.680 | 7.826 | **0.000746** |
|  | treatment x origin | 0.000441 | 0.000441 | 1 | 87.482 | 0.0326 | 0.857 |
|  | timepoint x treatment x origin | 0.0918 | 0.0459 | 2 | 88.212 | 3.393 | **0.0380** |
|  | Random effects | npar | logLik | AIC | LRT | Df | p-value |
|  | reduced | 14 | 60.018 | -92.0 |  |  |  |
|  | full | 13 | 59.489 | -93.0 | 1.0575 | 1 | 0.304 |
| ***Porites* sp.** | | | | | | | |
| **P_G day_/R_day_** | |  |  |  |  |  |  |
|  | GLMM | SS | MS | Chisq | Df | F statistic | p-value |
|  | timepoint | 0.0158 | 0.00789 | 0.862 | 2 | 0.291 | 0.650 |
|  | treatment | 0.00574 | 0.00574 | 0 | 1 | 0.212 | 1.000 |
|  | origin | 0.0266 | 0.0266 | 5.110 | 1 | 0.982 | **0.0238** |
|  | timepoint x treatment | 0.0487 | 0.0244 | 11.184 | 2 | 0.898 | **0.00373** |
|  | timepoint x origin | 0.368 | 0.184 | 1.283 | 2 | 6.779 | 0.527 |
|  | treatment x origin | 0.331 | 0.331 | 0 | 1 | 12.215 | 0.999 |
|  | timepoint x treatment x origin | 0.279 | 0.140 | 10.387 | 2 | 5.151 | **0.00555** |
|  | Random effects | npar | logLik | AIC | Chisq | Df | p-value |
|  | reduced | 13 | 9.932 | 6.137 |  |  |  |
|  | full | 14 | 13.593 | 0.814 | 7.323 | 1 | **0.00681** |
| **Chlorophyll a (µg cm^-2^)** | |  |  |  |  |  |  |
|  | LMM | SS | MS | Num df | Den df | F statistic | p-value |
|  | timepoint | 13.237 | 6.619 | 2 | 74.185 | 19.428 | **<0.0001** |
|  | treatment | 2.895 | 2.895 | 1 | 73.652 | 8.497 | **0.00471** |
|  | origin | 0.765 | 0.765 | 1 | 24.338 | 2.245 | 0.147 |
|  | timepoint x treatment | 9.389 | 4.694 | 2 | 72.294 | 13.780 | **<0.0001** |
|  | timepoint x origin | 5.509 | 2.755 | 2 | 74.185 | 8.085 | **0.000666** |
|  | treatment x origin | 1.201 | 1.201 | 1 | 73.652 | 3.525 | 0.0644 |
|  | timepoint x treatment x origin | 1.232 | 0.616 | 2 | 72.294 | 1.808 | 0.171 |
|  | Random effects | npar | logLik | AIC | LRT | Df | p-value |
|  | reduced | 14 | -90.224 | 208.5 |  |  |  |
|  | full | 13 | -92.696 | 211.4 | 4.944 | 1 | **0.0262** |
| **Symbiont density (x10^6^ cells cm^-2^)** | |  |  |  |  |  |  |
|  | LMM | SS | MS | Num df | Den df | F statistic | p-value |
|  | timepoint | 11.832 | 5.916 | 2 | 76.943 | 14.935 | **<0.0001** |
|  | treatment | 1.015 | 1.015 | 1 | 74.872 | 2.563 | 0.114 |
|  | origin | 1.047 | 1.047 | 1 | 23.918 | 2.643 | 0.117 |
|  | timepoint x treatment | 1.327 | 0.663 | 2 | 73.563 | 1.675 | 0.194 |
|  | timepoint x origin | 0.236 | 0.118 | 2 | 76.943 | 0.297 | 0.744 |
|  | treatment x origin | 0.295 | 0.295 | 1 | 74.872 | 0.745 | 0.391 |
|  | timepoint x treatment x origin | 0.238 | 0.119 | 2 | 73.563 | 0.300 | 0.741 |
|  | Random effects | npar | logLik | AIC | LRT | Df | p-value |
|  | reduced | 14 | -96.645 | 221.3 |  |  |  |
|  | full | 13 | -97.962 | 221.9 | 2.634 | 1 | 0.105 |
| **Tissue Biomass (mg cm^-2^)** | |  |  |  |  |  |  |
|  | LMM | SS | MS | Num df | Den df | F statistic | p-value |
|  | timepoint | 660.38 | 330.19 | 2 | 74.180 | 17.657 | **<0.0001** |
|  | treatment | 143.30 | 143.30 | 1 | 70.669 | 7.663 | **0.00719** |
|  | origin | 576.43 | 576.43 | 1 | 20.943 | 30.824 | **<0.0001** |
|  | timepoint x treatment | 206.34 | 103.17 | 2 | 70.051 | 5.517 | **0.00596** |
|  | timepoint x origin | 104.98 | 52.49 | 2 | 74.180 | 2.807 | 0.0668 |
|  | treatment x origin | 10.17 | 10.17 | 1 | 70.669 | 0.544 | 0.463 |
|  | timepoint x treatment x origin | 88.01 | 44.00 | 2 | 70.051 | 2.353 | 0.103 |
|  | Random effects | npar | logLik | AIC | LRT | Df | p-value |
|  | reduced | 14 | -248.44 | 524.9 |  |  |  |
|  | full | 13 | -248.75 | 523.5 | 0.612 | 1 | 0.434 |
| **Calcification (mg cm^-2^ d^-1^)** | |  |  |  |  |  |  |
|  | LMM | SS | MS | Num df | Den df | F statistic | p-value |
|  | timepoint | 0.0301 | 0.0301 | 1 | 46.084 | 0.624 | 0.434 |
|  | treatment | 1.276 | 1.276 | 1 | 46.156 | 26.446 | **<0.0001** |
|  | origin | 0.693 | 0.693 | 1 | 18.460 | 14.369 | **0.00129** |
|  | timepoint x treatment | 0.0236 | 0.0236 | 1 | 46.084 | 0.489 | 0.488 |
|  | timepoint x origin | 0.00662 | 0.00662 | 1 | 46.084 | 0.137 | 0.713 |
|  | treatment x origin | 0.68 | 0.688 | 1 | 46.156 | 14.260 | **0.000454** |
|  | timepoint x treatment x origin | 0.713 | 0.713 | 1 | 46.084 | 14.766 | **0.000371** |
|  | Random effects | npar | logLik | AIC | LRT | Df | p-value |
|  | reduced | 10 | -19.3 | 58.6 |  |  |  |
|  | full | 9 | -33.0 | 84.0 | 27.363 | 1 | **<0.0001** |
| **P_GROSS_ (μmol O2 cm^-2^ h^-1^)** | |  |  |  |  |  |  |
|  | LMM | SS | MS | Num df | Den df | F statistic | p-value |
|  | timepoint | 3.0355 | 1.518 | 2 | 85.752 | 10.324 | <0.0001 |
|  | treatment | 0.0528 | 0.0528 | 1 | 85.801 | 0.359 | 0.550 |
|  | origin | 0.0438 | 0.0438 | 1 | 18.239 | 0.298 | 0.592 |
|  | timepoint x treatment | 0.4353 | 0.218 | 2 | 85.752 | 1.481 | 0.233 |
|  | timepoint x origin | 10.114 | 5.0569 | 2 | 85.752 | 34.397 | **<0.0001** |
|  | treatment x origin | 0.591 | 0.590 | 1 | 85.801 | 4.0140 | **0.0483** |
|  | timepoint x treatment x origin | 0.308 | 0.1542 | 2 | 85.752 | 1.0472 | 0.356 |
|  | Random effects | npar | logLik | AIC | LRT | Df | p-value |
|  | reduced | 14 | -70.32 | 168.6 |  |  |  |
|  | full | 13 | -75.33 | 176.7 | 10.032 | 1 | **0.00154** |
| **R_DARK_ (μmol O2 cm^-2^ h^-1^)** | |  |  |  |  |  |  |
|  | GLMM | SS | MS | Chisq | Df | F statistic | p-value |
|  | timepoint | 0.640 | 0.320 | 15.093 | 2 | 6.833 | **0.000528** |
|  | treatment | 0.0112 | 0.0112 | 0 | 1 | 0.240 | 1.000 |
|  | origin | 0.0347 | 0.0347 | 1.822 | 1 | 0.742 | 0.177 |
|  | timepoint x treatment | 0.634 | 0.317 | 10.428 | 2 | 6.77 | **0.00544** |
|  | timepoint x origin | 1.205 | 0.603 | 16.868 | 2 | 12.874 | **0.000217** |
|  | treatment x origin | 0.0885 | 0.0885 | 0 | 1 | 1.89 | 1.000 |
|  | timepoint x treatment x origin | 0.0560 | 0.0280 | 1.300 | 2 | 0.598 | 0.522 |
|  | Random effects | npar | logLik | AIC | Chisq | Df | p-value |
|  | reduced | 13 | 71.59 | -117.2 |  |  |  |
|  | full | 14 | 74.38 | -120.8 | 5.57 | 1 | 0.0182 |
| **LEDR (μmol O2 cm^-2^ h^-1^)** | |  |  |  |  |  |  |
|  | LMM | SS | MS | Num df | Den df | F statistic | p-value |
|  | timepoint | 1.906 | 0.953 | 2 | 86.289 | 30.493 | **<0.0001** |
|  | treatment | 0.0601 | 0.0601 | 1 | 86.354 | 1.923 | 0.170 |
|  | origin | 0.0218 | 0.0218 | 1 | 18.571 | 0.697 | 0.414 |
|  | timepoint x treatment | 0.307 | 0.153 | 2 | 86.289 | 4.908 | **0.00957** |
|  | timepoint x origin | 1.127 | 0.564 | 2 | 86.289 | 18.0267 | **<0.0001** |
|  | treatment x origin | 0.00422 | 0.00422 | 1 | 86.354 | 0.135 | 0.714 |
|  | timepoint x treatment x origin | 0.0717 | 0.0358 | 2 | 86.289 | 1.146 | 0.323 |
|  | Random effects | npar | logLik | AIC | LRT | Df | p-value |
|  | reduced | 14 | 12.59 | 2.82 |  |  |  |
|  | full | 13 | 10.47 | 5.06 | 4.24 | 1 | **0.0395** |

**Table S5**

Summary of estimated marginal means (*post-hoc* tests) of on daily gross photosynthesis: daily respiration (P_G day_/R_day_), chlorophyll a concentrations, symbiont density, tissue biomass, calcification rates, gross photosynthesis (P_G_), dark respiration (R_dark_), and light-enhanced dark respiration (LEDR) for significant main effects or interactions of timepoint, origin, and treatment for *Siderastrea siderea* and branching *Porites* sp. Emmean = estimated marginal mean, SE = standard error, df = degrees of freedom, Lower.CL = lower confidence limit, Upper.CL = upper confidence limits, Group = different letters indicate statistically different groups.

| ***Siderastrea siderea*** | | | | | | | | | |
| --- | --- | --- | --- | --- | --- | --- | --- | --- | --- |
| **P_G day_/R_day_** | | | |  |  |  |  |  |  |
|  | timepoint | origin | treatment | emmeans | SE | df | Lower.CL | Upper.CL | Group |
|  | T0 | Reef | Native | 1.363 | 0.0618 | 95.7 | 1.241 | 1.49 | C |
|  |  |  | Transplant | 1.363 | 0.0618 | 95.7 | 1.241 | 1.49 | C |
|  |  | Bay | Native | 1.049 | 0.0618 | 95.7 | 0.927 | 1.17 | AB |
|  |  |  | Transplant | 1.049 | 0.0618 | 95.7 | 0.927 | 1.17 | AB |
|  | T4 | Reef | Native | 1.162 | 0.0618 | 95.7 | 1.039 | 1.28 | ABC |
|  |  |  | Transplant | 0.918 | 0.0618 | 95.7 | 0.796 | 1.04 | A |
|  |  | Bay | Native | 1.179 | 0.0650 | 98.0 | 1.050 | 1.31 | ABC |
|  |  |  | Transplant | 1.337 | 0.0650 | 98.2 | 1.208 | 1.47 | C |
|  | T12 | Reef | Native | 1.278 | 0.0618 | 95.7 | 1.155 | 1.40 | BC |
|  |  |  | Transplant | 1.030 | 0.0618 | 95.7 | 0.907 | 1.15 | AB |
|  |  | Bay | Native | 1.284 | 0.0618 | 95.7 | 1.162 | 1.41 | BC |
|  |  |  | Transplant | 1.353 | 0.0650 | 98.0 | 1.224 | 1.48 | C |
| **Chlorophyll a (µg cm^-2^)** | | | |  |  |  |  |  |  |
|  | timepoint | origin |  | emmeans | SE | df | Lower.CL | Upper.CL | Group |
|  | T0 | Reef |  | 1.188 | 0.0265 | 41.9 | 1.135 | 1.242 | D |
|  |  | Bay |  | 1.065 | 0.0279 | 45.0 | 1.008 | 1.121 | C |
|  | T4 | Reef |  | 0.989 | 0.0326 | 62.8 | 0.923 | 1.054 | BC |
|  |  | Bay |  | 0.896 | 0.0356 | 71.5 | 0.825 | 0.967 | AB |
|  | T12 | Reef |  | 0.835 | 0.0278 | 46.6 | 0.779 | 0.891 | A |
|  |  | Bay |  | 0.917 | 0.0265 | 41.9 | 0.863 | 0.970 | AB |
| **Symbiont density (x10^6^ cells cm^-2^)** | | | |  |  |  |  |  |  |
|  | timepoint |  |  | emmeans | SE | df | Lower.CL | Upper.CL | Group |
|  | T0 |  |  | 2.66 | 0.120 | 32.1 | 2.41 | 2.90 | B |
|  | T4 |  |  | 1.69 | 0.138 | 48.7 | 1.41 | 1.97 | A |
|  | T12 |  |  | 1.55 | 0.121 | 33.9 | 1.30 | 1.79 | A |
| **Biomass (mg cm^-2^)** | | | |  |  |  |  |  |  |
|  | timepoint | origin | treatment | emmeans | SE | df | Lower.CL | Upper.CL | Group |
|  | T0 | Reef | Native | 63.0 | 6.40 | 80.8 | 50.3 | 75.7 | A |
|  |  |  | Transplant | 63.0 | 6.40 | 80.8 | 50.3 | 75.7 | A |
|  |  | Bay | Native | 82.8 | 6.75 | 81.8 | 69.4 | 96.2 | ABC |
|  |  |  | Transplant | 82.8 | 6.75 | 81.8 | 69.4 | 96.2 | ABC |
|  | T4 | Reef | Native | 74.8 | 8.23 | 85.5 | 58.4 | 91.1 | ABC |
|  |  |  | Transplant | 80.3 | 8.24 | 85.5 | 63.9 | 96.7 | ABC |
|  |  | Bay | Native | 81.2 | 9.01 | 85.9 | 63.3 | 99.1 | ABC |
|  |  |  | Transplant | 102.5 | 24 | 85.4 | 86.1 | 118.9 | BC |
|  | T12 | Reef | Native | 74.9 | 6.40 | 80.8 | 62.2 | 87.7 | AB |
|  |  |  | Transplant | 86.0 | 7.15 | 83.5 | 71.8 | 100.2 | ABC |
|  |  | Bay | Native | 107.9 | 6.74 | 82.3 | 94.5 | 121.3 | C |
|  |  |  | Transplant | 85.1 | 6.40 | 80.8 | 72.3 | 97.8 | ABC |
| **Calcification (mg cm^-2^ d^-1^)** | | | |  |  |  |  |  |  |
|  |  | origin | treatment | emmeans | SE | df | Lower.CL | Upper.CL | Group |
|  |  | Reef | Native | 1.43 | 0.0964 | 30.5 | 1.236 | 1.63 | B |
|  |  |  | Transplant | 1.57 | 0.0964 | 30.5 | 1.375 | 1.77 | B |
|  |  | Bay | Native | 1.42 | 0.0981 | 32.0 | 1.215 | 1.62 | B |
|  |  |  | Transplant | 1.04 | 0.0981 | 32.0 | 0.838 | 1.24 | A |
| **P_GROSS_ (μmol O2 cm^-2^ h^-1^)** | | | |  |  |  |  |  |  |
|  | timepoint | origin |  | emmeans | SE | df | Lower.CL | Upper.CL | Group |
|  | T0 | Reef |  | 1.52 | 0.0469 | 55.5 | 1.43 | 1.61 | CD |
|  |  | Bay |  | 1.33 | 0.0469 | 55.5 | 1.24 | 1.42 | BC |
|  | T4 | Reef |  | 1.27 | 0.0469 | 55.5 | 1.18 | 1.37 | AB |
|  |  | Bay |  | 1.51 | 0.0489 | 60.9 | 1.42 | 1.61 | D |
|  | T12 | Reef |  | 1.12 | 0.0469 | 55.5 | 1.03 | 1.22 | A |
|  |  | Bay |  | 1.42 | 0.0480 | 58.1 | 1.33 | 1.52 | BCD |
|  |  | origin | treatment | emmeans | SE | df | Lower.CL | Upper.CL | Group |
|  |  | Reef | Native | 1.36 | 0.0409 | 35.9 | 1.28 | 1.45 | AB |
|  |  |  | Transplant | 1.25 | 0.0409 | 35.9 | 1.16 | 1.33 | A |
|  |  | Bay | Native | 1.41 | 0.0414 | 37.3 | 1.33 | 1.49 | B |
|  |  |  | Transplant | 1.43 | 0.0419 | 38.7 | 1.35 | 1.52 | B |
| **R_DARK_ (μmol O2 cm^-2^ h^-1^)** | | | |  |  |  |  |  |  |
|  | timepoint |  |  | emmeans | SE | df | Lower.CL | Upper.CL | Group |
|  | T0 |  |  | -0.140 | 0.0494 | inf | -0.237 | -0.0428 | B |
|  | T4 |  |  | -0.207 | 0.0502 | inf | -0.306 | -0.109 | B |
|  | T12 |  |  | -0.400 | 0.0501 | inf | -0.498 | -0.302 | A |
| **LEDR (μmol O2 cm^-2^ h^-1^)** | | | |  |  |  |  |  |  |
|  | timepoint | origin | treatment | emmeans | SE | df | Lower.CL | Upper.CL | Group |
|  | T0 | Reef | Native | -0.0140 | 0.0381 | 103 | -0.0896 | 0.0616 | C |
|  |  |  | Transplant | -0.0140 | 0.0381 | 103 | -0.0896 | 0.0616 | C |
|  |  | Bay | Native | -0.0464 | 0.0381 | 103 | -0.1220 | 0.0292 | BC |
|  |  |  | Transplant | -0.0464 | 0.0381 | 103 | -0.1220 | 0.0292 | BC |
|  | T4 | Reef | Native | -0.0530 | 0.0381 | 103 | -0.129 | 0.0266 | BC |
|  |  |  | Transplant | -0.185 | 0.0381 | 103 | -0.260 | -0.109 | ABC |
|  |  | Bay | Native | -0.0520 | 0.0402 | 103 | -0.132 | 0.0277 | BC |
|  |  |  | Transplant | -0.0337 | 0.0402 | 103 | -0.113 | 0.0460 | BC |
|  | T12 | Reef | Native | -0.275 | 0.0381 | 103 | -0.351 | -0.200 | A |
|  |  |  | Transplant | -0.210 | 0.0381 | 103 | -0.285 | -0.134 | AB |
|  |  | Bay | Native | -0.0371 | 0.0381 | 103 | -0.113 | 0.0385 | BC |
|  |  |  | Transplant | -0.0984 | 0.0402 | 103 | -0.178 | -0.0186 | ABC |
| ***Porites* sp.** | | | | | | | | | |
| **P_G day_/R_day_** | | | |  |  |  |  |  |  |
|  | timepoint | origin | treatment | emmeans | SE | df | Lower.CL | Upper.CL | Group |
|  | T0 | Reef | Native | 1.381 | 0.0820 | Inf | 1.221 | 1.54 | BC |
|  |  |  | Transplant | 1.381 | 0.0820 | Inf | 1.221 | 1.54 | BC |
|  |  | Bay | Native | 1.130 | 0.0717 | Inf | 0.989 | 1.27 | AB |
|  |  |  | Transplant | 1.130 | 0.0718 | Inf | 0.989 | 1.27 | AB |
|  | T4 | Reef | Native | 1.304 | 0.0785 | Inf | 1.150 | 1.46 | BC |
|  |  |  | Transplant | 1.171 | 0.0751 | Inf | 1.024 | 1.32 | ABC |
|  |  | Bay | Native | 1.206 | 0.0748 | Inf | 1.059 | 1.35 | ABC |
|  |  |  | Transplant | 1.446 | 0.0852 | Inf | 1.279 | 1.61 | C |
|  | T12 | Reef | Native | 1.371 | 0.0814 | Inf | 1.212 | 1.53 | BC |
|  |  |  | Transplant | 0.943 | 0.0794 | Inf | 0.787 | 1.10 | A |
|  |  | Bay | Native | 1.182 | 0.0767 | Inf | 1.031 | 1.33 | ABC |
|  |  |  | Transplant | 1.348 | 0.0810 | Inf | 1.189 | 1.51 | BC |
| **Chlorophyll a (µg cm^-2^)** | | | |  |  |  |  |  |  |
|  | timepoint | origin |  | emmeans | SE | df | Lower.CL | Upper.CL | Group |
|  | T0 | Reef |  | 4.53 | 0.159 | 41.4 | 4.21 | 4.85 | B |
|  |  | Bay |  | 4.28 | 0.167 | 44.9 | 3.94 | 4.62 | B |
|  | T4 | Reef |  | 2.92 | 0.205 | 65.3 | 2.51 | 3.33 | A |
|  |  | Bay |  | 3.92 | 0.197 | 61.2 | 3.53 | 4.32 | B |
|  | T12 | Reef |  | 4.09 | 0.183 | 54.6 | 3.73 | 4.46 | B |
|  |  | Bay |  | 4.12 | 0.161 | 71.9 | 3.80 | 4.44 | B |
|  | timepoint |  | treatment | emmeans | SE | df | Lower.CL | Upper.CL | Group |
|  | T0 |  | Native | 4.41 | 0.149 | 73.8 | 4.11 | 4.70 | B |
|  |  |  | Transplant | 4.41 | 0.149 | 73.8 | 4.11 | 4.70 | B |
|  | T4 |  | Native | 4.11 | 0.185 | 80.2 | 3.74 | 4.48 | B |
|  |  |  | Transplant | 2.73 | 0.194 | 80.7 | 2.34 | 3.12 | A |
|  | T12 |  | Native | 3.98 | 0.164 | 73.2 | 3.65 | 4.31 | B |
|  |  |  | Transplant | 4.23 | 0.167 | 77.8 | 3.90 | 4.57 | B |
| **Symbiont density (x10^6^ cells cm^-2^)** | | | |  |  |  |  |  |  |
|  | timepoint |  |  | emmeans | SE | df | Lower.CL | Upper.CL | Group |
|  | T0 |  |  | 2.15 | 0.119 | 46.7 | 1.91 | 2.39 | A |
|  | T4 |  |  | 1.81 | 0.147 | 68.1 | 1.51 | 2.10 | A |
|  | T12 |  |  | 2.74 | 0.122 | 67.2 | 2.50 | 2.99 | B |
| **Biomass (mg cm^-2^)** | | | |  |  |  |  |  |  |
|  | timepoint |  | treatment | emmeans | SE | df | Lower.CL | Upper.CL | Group |
|  | T0 |  | Native | 19.1 | 1.01 | 78.6 | 17.1 | 21.1 | A |
|  |  |  | Transplant | 19.1 | 1.01 | 78.6 | 17.1 | 21.1 | A |
|  | T4 |  | Native | 21.1 | 1.31 | 81.0 | 18.5 | 23.7 | A |
|  |  |  | Transplant | 22.0 | 1.37 | 81.0 | 19.3 | 24.8 | A |
|  | T12 |  | Native | 22.1 | 1.07 | 78.8 | 19.9 | 24.2 | A |
|  |  |  | Transplant | 29.0 | 1.32 | 81.0 | 26.3 | 31.6 | B |
| **Calcification (mg cm^-2^ d^-1^)** | | | |  |  |  |  |  |  |
|  | timepoint | origin | treatment | emmeans | SE | df | Lower.CL | Upper.CL | Group |
|  | T4 | Reef | Native | 2.16 | 0.111 | 32.3 | 1.932 | 2.39 | C |
|  |  |  | Transplant | 1.42 | 0.119 | 38.2 | 1.181 | 1.66 | A |
|  |  | Bay | Native | 1.21 | 0.111 | 32.3 | 0.983 | 1.44 | A |
|  |  |  | Transplant | 1.31 | 0.111 | 32.3 | 1.080 | 1.53 | A |
|  | T12 | Reef | Native | 1.85 | 0.111 | 32.3 | 1.619 | 2.07 | B |
|  |  |  | Transplant | 1.61 | 0.137 | 51.4 | 1.331 | 1.88 | AB |
|  |  | Bay | Native | 1.36 | 0.118 | 38.1 | 1.119 | 1.60 | AB |
|  |  |  | Transplant | 1.11 | 0.111 | 32.3 | 0.885 | 1.34 | A |
| **P_GROSS_ (μmol O2 cm^-2^ h^-1^)** | | | |  |  |  |  |  |  |
|  | timepoint | origin |  | emmeans | SE | df | Lower.CL | Upper.CL | Group |
|  | T0 | Reef |  | 2.29 | 0.110 | 45.1 | 2.07 | 2.51 | C |
|  |  | Bay |  | 1.43 | 0.110 | 45.1 | 1.21 | 1.65 | AB |
|  | T4 | Reef |  | 1.22 | 0.110 | 45.1 | 1.00 | 1.44 | A |
|  |  | Bay |  | 1.75 | 0.110 | 45.1 | 1.53 | 1.97 | B |
|  | T12 | Reef |  | 1.51 | 0.122 | 58.7 | 1.27 | 1.75 | AB |
|  |  | Bay |  | 1.65 | 0.112 | 47.5 | 1.42 | 1.87 | AB |
|  |  | origin | treatment | emmeans | SE | df | Lower.CL | Upper.CL | Group |
|  |  | Reef | Native | 1.73 | 0.0982 | 30.9 | 1.53 | 1.93 | A |
|  |  |  | Transplant | 1.62 | 0.104 | 37.1 | 1.41 | 1.84 | A |
|  |  | Bay | Native | 1.51 | 0.0992 | 31.7 | 1.31 | 1.72 | A |
|  |  |  | Transplant | 1.70 | 0.0982 | 30.6 | 1.50 | 1.90 | A |
| **R_DARK_ (μmol O2 cm^-2^ h^-1^)** | | | |  |  |  |  |  |  |
|  | timepoint | origin |  | emmeans | SE | df | Lower.CL | Upper.CL | Group |
|  | T0 | Reef |  | -0.399 | 0.0540 | 103 | -0.506 | -0.292 | B |
|  |  | Bay |  | -0.541 | 0.0540 | 103 | -0.648 | -0.434 | B |
|  | T4 | Reef |  | -0.789 | 0.0540 | 103 | -0.896 | -0.682 | A |
|  |  | Bay |  | -0.478 | 0.0540 | 103 | -0.585 | -0.371 | B |
|  | T12 | Reef |  | -0.425 | 0.0623 | 103 | -0.548 | -0.301 | B |
|  |  | Bay |  | -0.509 | 0.0555 | 103 | -0.619 | -0.399 | B |
|  | timepoint |  | treatment | emmeans | SE | df | Lower.CL | Upper.CL | Group |
|  | T0 |  | Native | -0.470 | 0.0540 | 103 | -0.577 | -0.363 | B |
|  |  |  | Transplant | -0.470 | 0.0540 | 103 | -0.577 | -0.363 | B |
|  | T4 |  | Native | -0.566 | 0.0540 | 103 | -0.673 | -0.459 | AB |
|  |  |  | Transplant | -0.701 | 0.0540 | 103 | -0.809 | -0.594 | A |
|  | T12 |  | Native | -0.587 | 0.0555 | 103 | -0.697 | -0.477 | AB |
|  |  |  | Transplant | -0.347 | 0.0623 | 103 | -0.470 | -0.223 | B |
| **LEDR (μmol O2 cm^-2^ h^-1^)** | | | |  |  |  |  |  |  |
|  | timepoint | origin |  | emmeans | SE | df | Lower.CL | Upper.CL | Group |
|  | T0 | Reef |  | -1.056 | 0.0459 | 56.7 | -1.148 | -0.964 | A |
|  |  | Bay |  | -0.756 | 0.0459 | 56.7 | -0.848 | -0.664 | B |
|  | T4 | Reef |  | -0.515 | 0.0459 | 56.7 | -0.607 | -0.423 | C |
|  |  | Bay |  | -0.683 | 0.0459 | 56.7 | -0.774 | -0.591 | BC |
|  | T12 | Reef |  | -0.776 | 0.0519 | 71.4 | -0.879 | -0.672 | B |
|  |  | Bay |  | -0.791 | 0.0470 | 59.5 | -0.885 | -0.697 | B |
|  | timepoint |  | treatment | emmeans | SE | df | Lower.CL | Upper.CL | Group |
|  | T0 |  | Native | -0.906 | 0.0428 | 93.2 | -0.991 | -0.821 | A |
|  |  |  | Transplant | -0.906 | 0.0428 | 93.2 | -0.991 | -0.821 | A |
|  | T4 |  | Native | -0.627 | 0.0428 | 93.2 | -0.712 | -0.542 | B |
|  |  |  | Transplant | -0.571 | 0.0428 | 93.2 | -0.656 | -0.486 | B |
|  | T12 |  | Native | -0.686 | 0.0440 | 94.7 | -0.773 | -0.598 | B |
|  |  |  | Transplant | -0.881 | 0.0492 | 99.2 | -0.978 | -0.783 | A |

**Table S6**

Results of linear mixed models (LMM) and generalized linear mixed models (GLMM) analyses to test for the main effects of treatment and origin, their interaction, and the random effect of genotype on calcification rates between T0 and T12 and tissue biomass at T12 for *Siderastrea siderea* and branching *Porites* sp. P-values ≤ 0.05 are highlighted in bold. SS = Sum of squares, MS = Mean squares, Num df = numerator degrees of freedom, Den df = denominator degrees of freedom, Chisq = Chi-squared statistic, npar = number of parameters, logLik = log-liklihood, LRT = likelihood ratio test, Df = degrees of freedom. For *post-hoc* test results, see Table S7.

| ***Siderastrea siderea*** | | | | | | | |
| --- | --- | --- | --- | --- | --- | --- | --- |
| **Calcification (mg cm^-2^ d^-1^)** | |  |  |  |  |  |  |
|  | LMM | SS | MS | Num df | Den df | F statistic | p-value |
|  | treatment | 0.335 | 0.335 | 1 | 17.347 | 3.662 | 0.0723 |
|  | origin | 0.123 | 0.123 | 1 | 17.564 | 1.351 | 0.261 |
|  | treatment x origin | 1.171 | 1.171 | 1 | 17.347 | 12.813 | **0.00225** |
|  | Random effects | npar | logLik | AIC | LRT | Df | p-value |
|  | reduced | 6 | -16.034 | 44.069 |  |  |  |
|  | full | 5 | -16.395 | 42.790 | 0.722 | 1 | 0.396 |
| **Tissue Biomass (mg cm^-2^)** | |  |  |  |  |  |  |
|  | LMM | SS | MS | Num df | Den df | F statistic | p-value |
|  | treatment | 354.18 | 354.18 | 1 | 33 | 0.7671 | 0.38745 |
|  | origin | 2442.82 | 2442.82 | 1 | 33 | 5.2907 | **0.0279** |
|  | treatment x origin | 2660.23 | 2660.23 | 1 | 33 | 5.7615 | **0.00218** |
|  | Random effects | npar | logLik | AIC | LRT | Df | p-value |
|  | reduced | 6 | -152.49 | 317.0 |  |  |  |
|  | full | 5 | -152.49 | 315.0 | -5.68 x 10^-14^ | 1 | 1 |
| ***Porites*** **sp.** | | | | | | | |
| **Calcification (mg cm^-2^ d^-1^)** | |  |  |  |  |  |  |
|  | LMM | SS | MS | Num df | Den df | F statistic | p-value |
|  | treatment | 0.525 | 0.525 | 1 | 14.743 | 6.153 | **0.0257** |
|  | origin | 0.631 | 0.631 | 1 | 19.472 | 7.394 | **0.0134** |
|  | treatment x origin | 0.00154 | 0.00154 | 1 | 14.743 | 0.0181 | 0.895 |
|  | Random effects | npar | logLik | AIC | LRT | Df | p-value |
|  | reduced | 6 | -17.670 | 47.340 |  |  |  |
|  | full | 5 | -19.413 | 48.827 | 3.487 | 1 | **0.0619** |
| **Tissue Biomass (mg cm^-2^)** | |  |  |  |  |  |  |
|  | LMM | SS | MS | Num df | Den df | F statistic | p-value |
|  | treatment | 298.843 | 298.843 | 1 | 18.938 | 27.4028 | **4.77 x 10^-5^** |
|  | origin | 26.080 | 26.080 | 1 | 18.340 | 2.3915 | 0.1391 |
|  | treatment x origin | 86.425 | 86.425 | 1 | 18.938 | 7.9249 | **0.01108** |
|  | Random effects | npar | logLik | AIC | LRT | Df | p-value |
|  | reduced | 6 | -76.437 | 164.88 |  |  |  |
|  | full | 5 | -76.687 | 163.38 | 0.50003 | 1 | **0.4795** |

Table S7

Summary of estimated marginal means (*post-hoc* tests) of calcification rates and tissue biomass during the final timepoint only for *Siderastrea siderea* and branching *Porites* sp. Emmean = estimated marginal mean, SE = standard error, df = degrees of freedom, Lower.CL = lower confidence limit, Upper.CL = upper confidence limits, Group = different letters indicate statistically different groups

| ***Siderastrea siderea*** | | | | | | | | | |
| --- | --- | --- | --- | --- | --- | --- | --- | --- | --- |
| **Calcification (mg cm^-2^ d^-1^)** | | | |  |  |  |  |  |  |
|  |  | origin | treatment | emmeans | SE | df | Lower.CL | Upper.CL | Group |
|  |  | Reef | Native | 1.46 | 0.107 | 33.6 | 1.238 | 1.67 | AB |
|  |  |  | Transplant | 1.62 | 0.107 | 33.6 | 1.400 | 1.84 | B |
|  |  | Bay | Native | 1.67 | 0.107 | 33.6 | 1.448 | 1.88 | B |
|  |  |  | Transplant | 1.13 | 0.114 | 34.1 | 0.901 | 1.36 | A |
| **Tissue biomass (mg cm^-2^)** | | | |  |  |  |  |  |  |
|  |  | origin | treatment | emmeans | SE | df | Lower.CL | Upper.CL | Group |
|  |  | Reef | Native | 18.9 | 1.30 | 22.9 | 16.2 | 21.6 | A |
|  |  |  | Transplant | 30.1 | 1.86 | 24.8 | 26.3 | 33.9 | B |
|  |  | Bay | Native | 25.2 | 1.40 | 24.9 | 22.3 | 28.1 | B |
|  |  |  | Transplant | 28.6 | 1.50 | 24.9 | 25.5 | 31.7 | B |
| ***Porites* sp.** | | | | | | | | | |
| **Calcification (mg cm^-2^ d^-1^)** | | | |  |  |  |  |  |  |
|  |  | origin |  | emmeans | SE | df | Lower.CL | Upper.CL | Group |
|  |  | Reef |  | 1.70 | 0.126 | 20.4 | 1.44 | 1.96 | B |
|  |  | Bay |  | 1.24 | 0.114 | 16.2 | 1.00 | 1.48 | A |
|  |  |  | treatment | emmeans | SE | df | Lower.CL | Upper.CL | Group |
|  |  |  | Native | 1.61 | 0.0955 | 26.4 | 1.41 | 1.81 | B |
|  |  |  | Transplant | 1.33 | 0.109 | 28.7 | 1.11 | 1.56 | A |
| **Tissue biomass (mg cm^-2^)** | | | |  |  |  |  |  |  |
|  |  | origin | treatment | emmeans | SE | df | Lower.CL | Upper.CL | Group |
|  |  | Reef | Native | 18.9 | 1.30 | 22.9 | 16.2 | 21.6 | A |
|  |  |  | Transplant | 30.1 | 1.86 | 24.8 | 26.3 | 33.9 | B |
|  |  | Bay | Native | 25.2 | 1.40 | 24.9 | 22.3 | 28.1 | B |
|  |  |  | Transplant | 28.6 | 1.50 | 24.9 | 25.5 | 31.7 | B |

**Table S8**

Results of (G)LMMs to assess effects of origin (bay or reef), treatment (native or transplant), and timepoint (T0, T4, T12) on chlorophyll a concentrations per cell, area-normalized chlorophyll c_2_ concentrations, and chlorophyll c_2_ concentrations per cell. (G)LMM = (Generalized) Linear mixed model, SS = Sum of squares, MS = Mean squares, Num df = numerator degrees of freedom, Den df = denominator degrees of freedom, Chisq = Chi-squared statistic, npar = number of parameters, logLik = log-liklihood, AIC = Akaike Information Criterion, LRT = likelihood ratio test, Df = degrees of freedom. *Post-hoc* results follow with summary of estimated marginal means. Emmean = estimated marginal mean, SE = standard error, df = degrees of freedom, Lower.CL = lower confidence limit, Upper.CL = upper confidence limits.

| ***Siderastrea siderea*** | | | | | | | | | | | | | | |
| --- | --- | --- | --- | --- | --- | --- | --- | --- | --- | --- | --- | --- | --- | --- |
| **Chlorophyll a (pg cell^-1^)** | | | |  | |  | |  | |  | |  | |  |
|  | | LMM | | SS | | MS | | Num df | | Den df | | F statistic | | p-value |
|  | | timepoint | | 0.0306 | | 0.015 | | 2 | | 67.502 | | 0.936 | | 0.397 |
|  |  | treatment | | 0.000 | | 0.000 | | 1 | | 68.105 | | 0.000 | | 0.990 |
|  |  | origin | | 0.110 | | 0.110 | | 1 | | 15.979 | | 6.719 | | **0.0196** |
|  |  | timepoint x treatment | | 0.000423 | | 0.000 | | 2 | | 66.725 | | 0.013 | | 0.987 |
|  |  | timepoint x origin | | 0.131 | | 0.066 | | 2 | | 67.502 | | 4.007 | | **0.0227** |
|  |  | treatment x origin | | 0.0961 | | 0.096 | | 1 | | 68.105 | | 5.874 | | **0.0180** |
|  |  | timepoint x treatment x origin | | 0.0717 | | 0.036 | | 2 | | 66.725 | | 2.191 | | 0.120 |
|  | | Random effects | | npar | | logLik | | AIC | | LRT | | Df | | p-value |
|  | | reduced | | 14 | | 34.408 | | -40.8 | |  | |  | |  |
|  | | full | | 13 | | 32.525 | | -39.1 | | 3.765 | | 1 | | 0.0523 |
| **Chlorophyll c_2_ (µg cm^-2^)** | | | |  | |  | |  | |  | |  | |  |
|  | | LMM | | SS | | MS | | Num df | | Den df | | F statistic | | p-value |
|  | | timepoint | | 3.169 | | 1.584 | | 2 | | 69.084 | | 41.930 | | **<0.0001** |
|  |  | treatment | | 0.109 | | 0.109 | | 1 | | 69.634 | | 2.890 | | 0.0936 |
|  |  | origin | | 0.306 | | 0.306 | | 1 | | 16.764 | | 8.103 | | **0.0113** |
|  |  | timepoint x treatment | | 0.106 | | 0.053 | | 2 | | 68.642 | | 1.404 | | 0.253 |
|  |  | timepoint x origin | | 1.839 | | 0.919 | | 2 | | 69.084 | | 24.333 | | **<0.0001** |
|  |  | treatment x origin | | 0.101 | | 0.101 | | 1 | | 69.634 | | 2.663 | | 0.107 |
|  |  | timepoint x treatment x origin | | 0.255 | | 0.128 | | 2 | | 68.642 | | 3.378 | | **0.0399** |
|  | | Random effects | | npar | | logLik | | AIC | | LRT | | Df | | p-value |
|  | | reduced | | 14 | | -2.272 | | 32.5 | |  | |  | |  |
|  | | full | | 13 | | -5.770 | | 37.5 | | 7.0 | | 1 | | **0.00817** |
| **Chlorophyll c_2_ (pg cell^-1^)** | | | |  | |  | |  | |  | |  | |  |
|  | | LMM | | SS | | MS | | Num df | | Den df | | F statistic | | p-value |
|  | | timepoint | | 0.00096 | | 0.00048 | | 2 | | 67.615 | | 0.0034 | | 0.997 |
|  |  | treatment | | 0.213 | | 0.213 | | 1 | | 67.728 | | 1.490 | | 0.227 |
|  |  | origin | | 1.0764 | | 1.0764 | | 1 | | 17.225 | | 7.531 | | **0.0137** |
|  |  | timepoint x treatment | | 0.202 | | 0.101 | | 2 | | 66.845 | | 0.706 | | 0.497 |
|  |  | timepoint x origin | | 2.557 | | 1.278 | | 2 | | 67.615 | | 8.945 | | **0.000358** |
|  |  | treatment x origin | | 0.979 | | 0.979 | | 1 | | 67.728 | | 6.852 | | **0.0109** |
|  |  | timepoint x treatment x origin | | 0.910 | | 0.455 | | 2 | | 66.845 | | 3.183 | | **0.0478** |
|  | | Random effects | | npar | | logLik | | AIC | | LRT | | Df | | p-value |
|  | | reduced | | 14 | | -60.49 | | 149.0 | |  | |  | |  |
|  | | full | | 13 | | -65.85 | | 157.7 | | 10.719 | | 1 | | **0.00106** |
| ***Porites* sp.** | | | | | | | | | | | | | | |
| **Chlorophyll a (pg cell^-1^)** | | | |  | |  | |  | |  | |  | |  |
|  | LMM | | SS | | MS | | Num df | | Den df | | F statistic | | p-value | |
|  | | timepoint | | 198.685 | | 99.343 | | 2 | | 79 | | 13.967 | | **<0.0001** |
|  |  | treatment | | 0.240 | | 0.240 | | 1 | | 79 | | 0.0338 | | 0.855 |
|  |  | origin | | 35.277 | | 35.277 | | 1 | | 79 | | 4.960 | | **0.0288** |
|  |  | timepoint x treatment | | 5.801 | | 2.901 | | 2 | | 79 | | 0.408 | | 0.666 |
|  |  | timepoint x origin | | 25.703 | | 12.852 | | 2 | | 79 | | 1.807 | | 0.171 |
|  |  | treatment x origin | | 4.356 | | 4.356 | | 1 | | 79 | | 0.612 | | 0.436 |
|  |  | timepoint x treatment x origin | | 2.520 | | 1.260 | | 2 | | 79 | | 0.177 | | 0.838 |
|  | | Random effects | | npar | | logLik | | AIC | | LRT | | Df | | p-value |
|  | | reduced | | 14 | | -201.59 | | 431.2 | |  | |  | |  |
|  | | full | | 13 | | -201.59 | | 429.2 | | <0.0001 | | 1 | | 1 |
| **Chlorophyll c_2_ (µg cm^-2^)** | | | |  | |  | |  | |  | |  | |  |
|  | | LMM | | SS | | MS | | Num df | | Den df | | F statistic | | p-value |
|  | | timepoint | | 47.937 | | 23.9687 | | 2 | | 73.817 | | 13.6508 | | **<0.0001** |
|  |  | treatment | | 2.317 | | 2.3173 | | 1 | | 73.465 | | 1.3197 | | 0.254 |
|  |  | origin | | 1.281 | | 1.2811 | | 1 | | 23.412 | | 0.7296 | | 0.402 |
|  |  | timepoint x treatment | | 23.950 | | 11.9750 | | 2 | | 72.093 | | 6.8200 | | 0.00194 |
|  |  | timepoint x origin | | 10.760 | | 5.3798 | | 2 | | 73.817 | | 3.0639 | | 0.0527 |
|  |  | treatment x origin | | 0.606 | | 0.6063 | | 1 | | 73.465 | | 0.3453 | | 0.559 |
|  |  | timepoint x treatment x origin | | 10.891 | | 5.4457 | | 2 | | 72.093 | | 3.1015 | | 0.0510 |
|  | | Random effects | | npar | | logLik | | AIC | | LRT | | Df | | p-value |
|  | | reduced | | 14 | | -158.27 | | 344.5 | |  | |  | |  |
|  | | full | | 13 | | -161.87 | | 349.7 | | 7.198 | | 1 | | **0.00730** |
| **Chlorophyll c_2_ (pg cell^-1^)** | | | |  | |  | |  | |  | |  | |  |
|  | | LMM | | SS | | MS | | Num df | | Den df | | F statistic | | p-value |
|  | | timepoint | | 12.853 | | 6.426 | | 2 | | 73.407 | | 12.601 | | **<0.0001** |
|  |  | treatment | | 0.228 | | 0.228 | | 1 | | 71.534 | | 0.447 | | 0.506 |
|  |  | origin | | 1.170 | | 1.170 | | 1 | | 22.161 | | 2.295 | | 0.144 |
|  |  | timepoint x treatment | | 0.691 | | 0.346 | | 2 | | 69.523 | | 0.678 | | 0.511 |
|  |  | timepoint x origin | | 3.921 | | 1.960 | | 2 | | 73.407 | | 3.844 | | **0.0258** |
|  |  | treatment x origin | | 0.107 | | 0.107 | | 1 | | 71.534 | | 0.210 | | 0.648 |
|  |  | timepoint x treatment x origin | | 1.0104 | | 0.505 | | 2 | | 69.523 | | 0.991 | | 0.377 |
|  | | Random effects | | npar | | logLik | | AIC | | LRT | | Df | | p-value |
|  | | reduced | | 14 | | -99.272 | | 226.5 | |  | |  | |  |
|  | | full | | 13 | | -99.411 | | 224.8 | | 0.278 | | 1 | | 0.598 |

**Table S9**

Summary of estimated marginal means (*post-hoc* tests) of chlorophyll a concentrations per cell, area-normalized chlorophyll c_2_ concentrations, and chlorophyll c_2_ concentrations per cell. Emmean = estimated marginal mean, SE = standard error, df = degrees of freedom, Lower.CL = lower confidence limit, Upper.CL = upper confidence limits, Group = different letters indicate statistically different groups.

| ***Siderastrea siderea*** | | | | | | | | | |
| --- | --- | --- | --- | --- | --- | --- | --- | --- | --- |
| **Chlorophyll a (pg cell^-1^)** | | | |  |  |  |  |  |  |
|  | timepoint | origin |  | emmeans | SE | df | Lower.CL | Upper.CL | Group |
|  | T0 | Reef |  | 0.809 | 0.0350 | 42.0 | 0.738 | 0.880 | B |
|  |  | Bay |  | 0.636 | 0.0368 | 45.0 | 0.562 | 0.710 | A |
|  | T4 | Reef |  | 0.815 | 0.0432 | 62.5 | 0.728 | 0.901 | B |
|  |  | Bay |  | 0.684 | 0.0473 | 70.5 | 0.590 | 0.778 | AB |
|  | T12 | Reef |  | 0.704 | 0.0367 | 46.7 | 0.630 | 0.778 | AB |
|  |  | Bay |  | 0.698 | 0.0367 | 46.8 | 0.624 | 0.771 | AB |
|  |  | origin | treatment | emmeans | SE | df | Lower.CL | Upper.CL | Group |
|  |  | Reef | Native | 0.809 | 0.0331 | 33.8 | 0.742 | 0.876 | B |
|  |  |  | Transplant | 0.743 | 0.0341 | 35.7 | 0.673 | 0.812 | AB |
|  |  | Bay | Native | 0.639 | 0.0361 | 41.7 | 0.566 | 0.712 | A |
|  |  |  | Transplant | 0.706 | 0.0344 | 36.6 | 0.637 | 0.776 | AB |
| **Chlorophyll c_2_ (µg cm^-2^)** | | | |  |  |  |  |  |  |
|  | timepoint | origin | treatment | emmeans | SE | df | Lower.CL | Upper.CL | Group |
|  | T0 | Reef | Native | 1.834 | 0.0714 | 68.0 | 1.692 | 1.980 | E |
|  |  |  | Transplant | 1.834 | 0.0714 | 68.0 | 1.692 | 1.980 | E |
|  |  | Bay | Native | 1.607 | 0.0750 | 71.2 | 1.458 | 1.760 | CDE |
|  |  |  | Transplant | 1.607 | 0.0750 | 71.2 | 1.458 | 1.760 | CDE |
|  | T4 | Reef | Native | 1.753 | 0.0896 | 82.2 | 1.574 | 1.930 | DE |
|  |  |  | Transplant | 1.454 | 0.0897 | 82.1 | 1.276 | 1.630 | ABCD |
|  |  | Bay | Native | 0.988 | 0.1081 | 85.8 | 0.773 | 1.200 | A |
|  |  |  | Transplant | 1.160 | 0.0897 | 81.9 | 0.982 | 1.340 | A |
|  | T12 | Reef | Native | 1.316 | 0.0714 | 68.0 | 1.174 | 1.460 | ABC |
|  |  |  | Transplant | 1.198 | 0.0788 | 75.4 | 1.041 | 1.350 | AB |
|  |  | Bay | Native | 1.537 | 0.0714 | 68.0 | 1.394 | 1.680 | BCDE |
|  |  |  | Transplant | 1.356 | 0.0714 | 68.0 | 1.213 | 1.500 | ABC |
| **Chlorophyll c_2_ (pg cell^-1^)** | | | |  |  |  |  |  |  |
|  | timepoint | origin | treatment | emmeans | SE | df | Lower.CL | Upper.CL | Group |
|  | T0 | Reef | Native | 1.457 | 0.147 | 58.8 | 1.164 | 1.75 | AB |
|  |  |  | Transplant | 1.457 | 0.147 | 58.8 | 1.164 | 1.75 | AB |
|  |  | Bay | Native | 1.054 | 0.153 | 62.5 | 0.747 | 1.36 | A |
|  |  |  | Transplant | 1.054 | 0.153 | 62.5 | 0.747 | 1.36 | A |
|  | T4 | Reef | Native | 1.985 | 0.181 | 76.6 | 1.625 | 2.35 | B |
|  |  |  | Transplant | 1.372 | 0.181 | 76.6 | 1.011 | 1.73 | AB |
|  |  | Bay | Native | 0.622 | 0.217 | 82.9 | 0.191 | 1.05 | A |
|  |  |  | Transplant | 1.055 | 0.181 | 76.2 | 0.693 | 1.42 | A |
|  | T12 | Reef | Native | 1.395 | 0.147 | 58.8 | 1.102 | 1.69 | AB |
|  |  |  | Transplant | 1.063 | 0.161 | 67.5 | 0.742 | 1.38 | A |
|  |  | Bay | Native | 1.316 | 0.147 | 58.8 | 1.022 | 1.61 | AB |
|  |  |  | Transplant | 1.227 | 0.161 | 67.5 | 0.907 | 1.55 | AB |
| ***Porites* sp.** | | | | | | | | | |
|  | **Chlorophyll a (pg cell^-1^)** | | |  |  |  |  |  |  |
|  | timepoint |  |  | emmeans | SE | df | Lower.CL | Upper.CL | Group |
|  | T0 |  |  | 9.97 | 0.447 | 58.9 | 9.08 | 10.87 | B |
|  | T4 |  |  | 8.06 | 0.565 | 70.2 | 6.94 | 9.19 | A |
|  | T12 |  |  | 6.53 | 0.485 | 72.5 | 5.56 | 7.50 | A |
|  |  | origin |  | emmeans | SE | df | Lower.CL | Upper.CL | Group |
|  |  | Reef |  | 8.83 | 0.416 | 16.1 | 7.95 | 9.71 | B |
|  |  | Bay |  | 7.55 | 0.400 | 25.2 | 6.73 | 8.37 | A |
|  | **Chlorophyll c_2_ (µg cm^-2^)** | | |  |  |  |  |  |  |
|  | timepoint |  |  | emmeans | SE | df | Lower.CL | Upper.CL | Group |
|  | T0 |  |  | 9.97 | 0.447 | 58.9 | 9.08 | 10.87 | B |
|  | T4 |  |  | 8.06 | 0.565 | 70.2 | 6.94 | 9.19 | A |
|  | T12 |  |  | 6.53 | 0.485 | 72.5 | 5.56 | 7.50 | A |
|  |  | origin |  | emmeans | SE | df | Lower.CL | Upper.CL | Group |
|  |  | Reef |  | 8.83 | 0.416 | 16.1 | 7.95 | 9.71 | B |
|  |  | Bay |  | 7.55 | 0.400 | 25.2 | 6.73 | 8.37 | A |
|  | **Chlorophyll c_2_ (pg cell^-1^)** | | |  |  |  |  |  |  |
|  | timepoint | origin |  | emmeans | SE | df | Lower.CL | Upper.CL | Group |
|  | T0 | Reef |  | 2.62 | 0.177 | 54.9 | 2.268 | 2.98 | B |
|  |  | Bay |  | 2.02 | 0.177 | 54.6 | 1.670 | 2.38 | AB |
|  | T4 | Reef |  | 2.00 | 0.226 | 70.3 | 1.549 | 2.45 | AB |
|  |  | Bay |  | 1.51 | 0.217 | 66.9 | 1.073 | 1.94 | A |
|  | T12 | Reef |  | 1.28 | 0.199 | 60.4 | 0.887 | 1.68 | A |
|  |  | Bay |  | 1.62 | 0.181 | 77.8 | 1.255 | 1.98 | A |

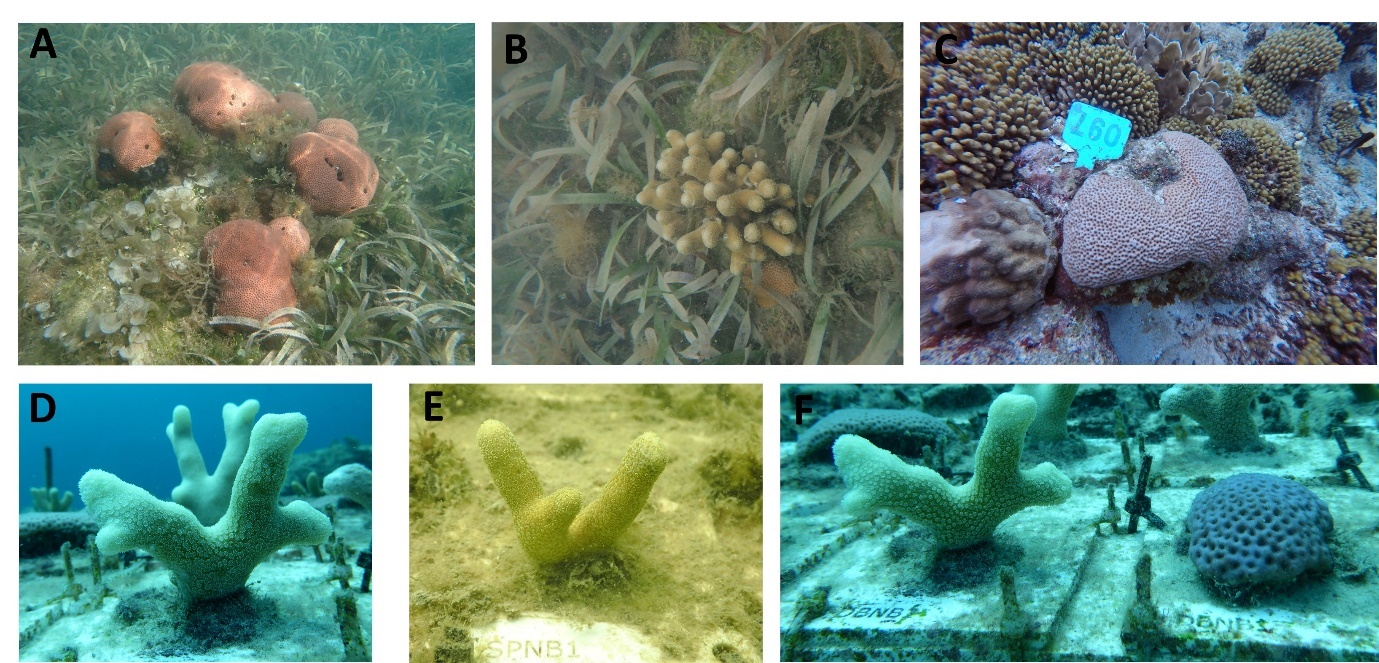

**Figure S1**

Photos from Spanish Water bay A-B) *Siderastrea siderea* colonies and isolated colony of branching *Porites* sp. growing among seagrass and C) reef-origin *S. siderea*. Photos comparing morphology of reef-origin *Porites* (D) and bay-origin *Porites* (E). Photo of reef-origin *Porites* and *S. siderea* on transplant structure at Carmabi House Reef (F).

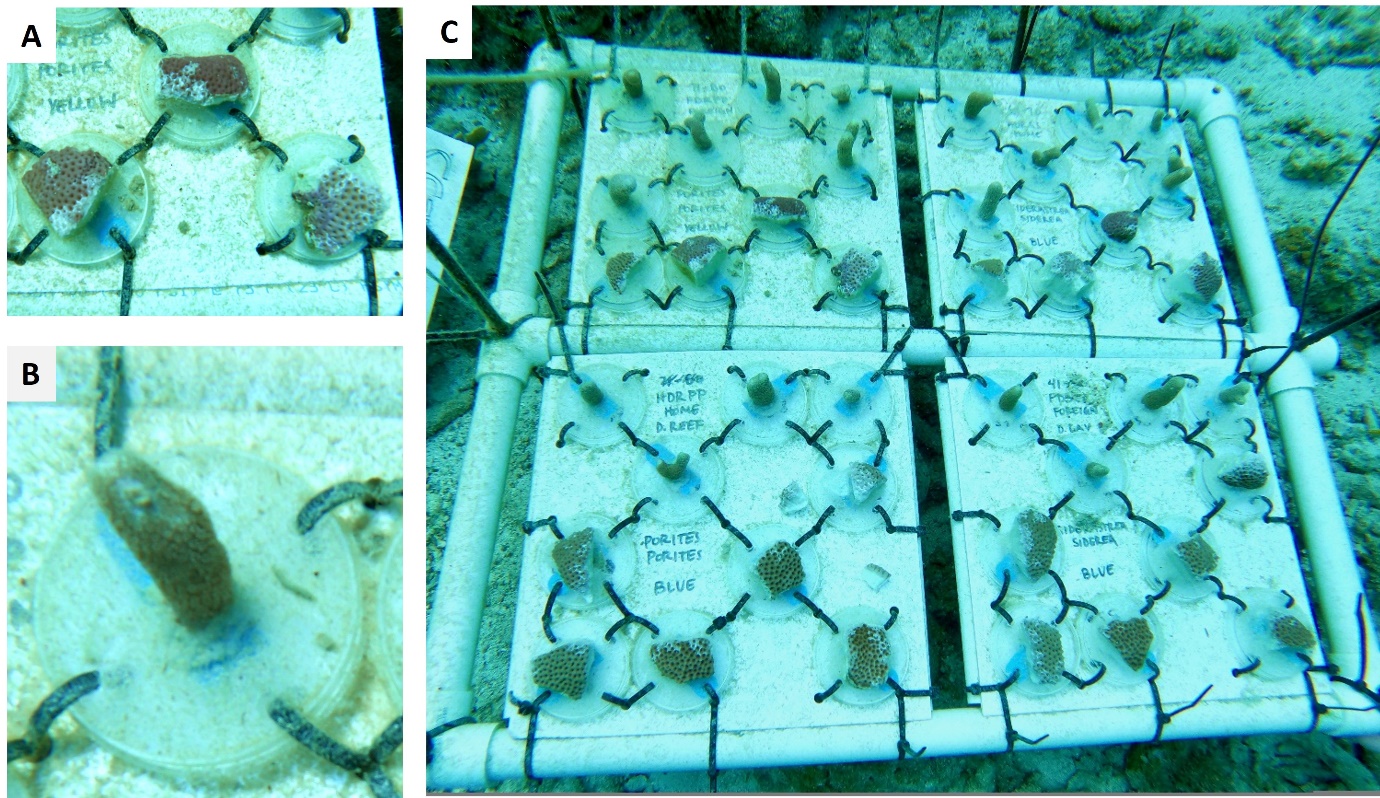

**Figure S2**

Photos illustrating the sudden partial mortality of A) *Siderastrea siderea* and B) branching *Porites* sp. observed during a separate pilot transplantation study that took place at Tugboat Reef prior to the start of the transplant experiment described here (see Methods for details). Panel C is a photo of the entire table structure.

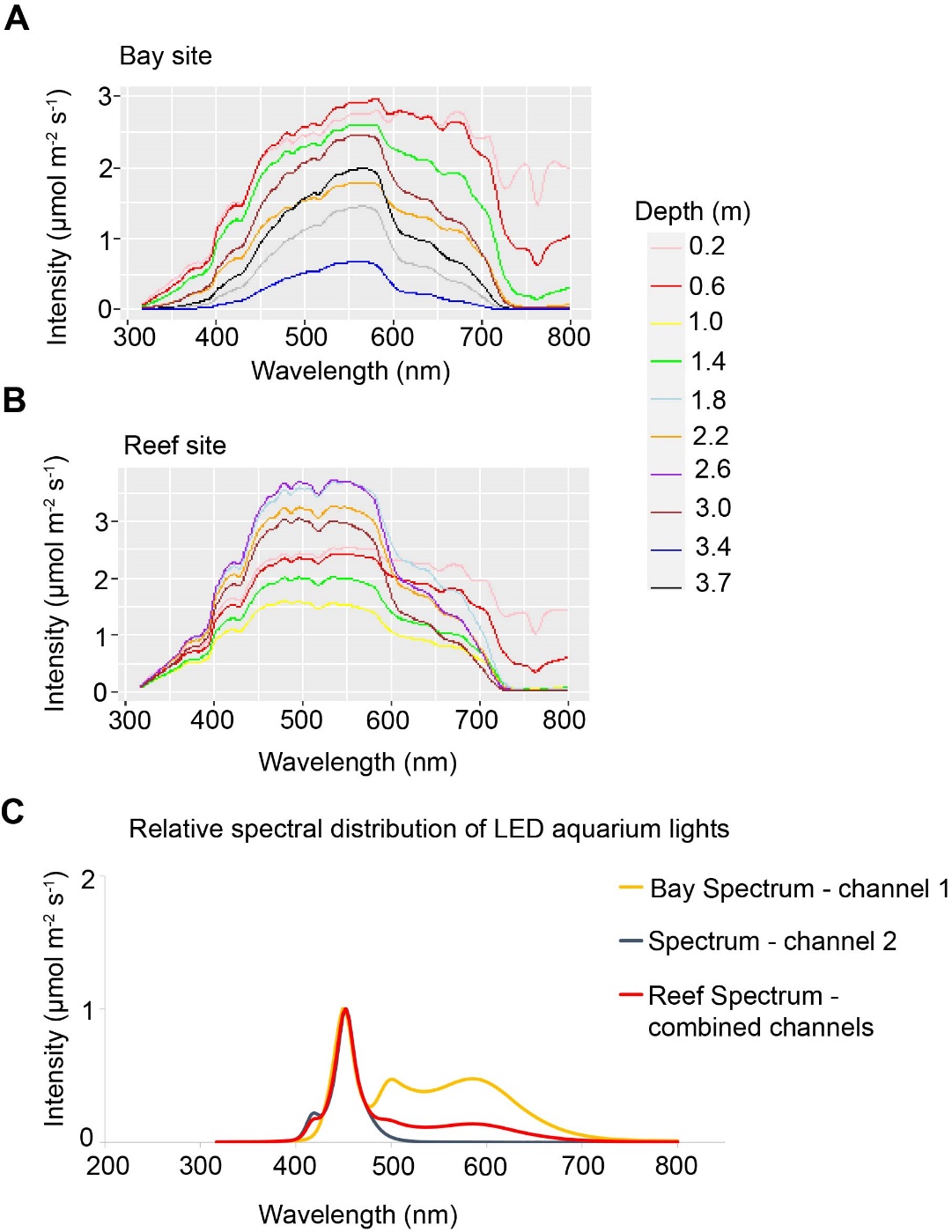

**Figure S3**

Light spectral profiles measured at a **A)** bay site and **B)** reef site across multiple depths. **C)** Light spectra of Phillips CoralCare Gen2 LED aquarium lights used for bay (yellow line) and reef (red line) during P:R incubations. Each channel represents the following settings : Channel 1 = “100% warm : 0% cool”, channel 2 = “0% warm : 100% cool”, and combined channels = “50% warm : 50% cool”. See Methods for more details.

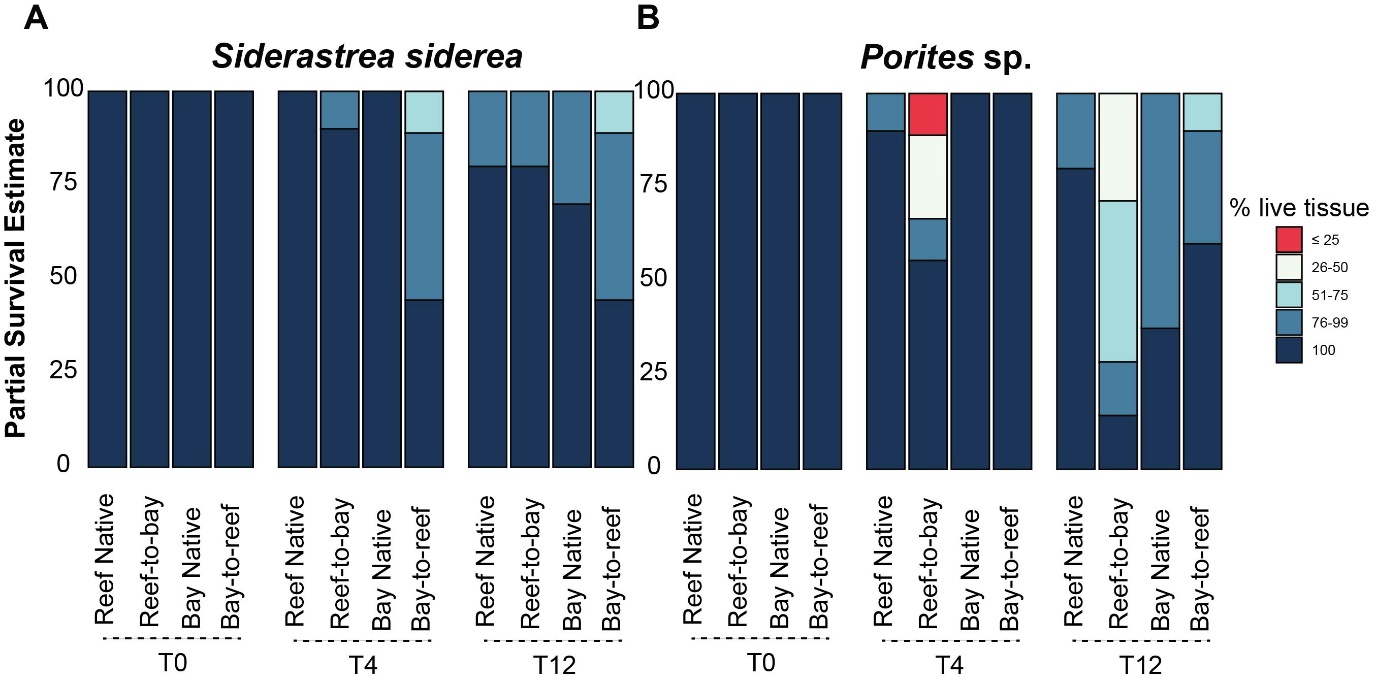

**Figure S4**

Partial survival estimates represented by percentage of living tissue for A) *Siderastrea siderea* and B) branching *Porites* sp. of all four transplant groups after 0, 4, and 12 months of transplantation (T0, T4, T12).

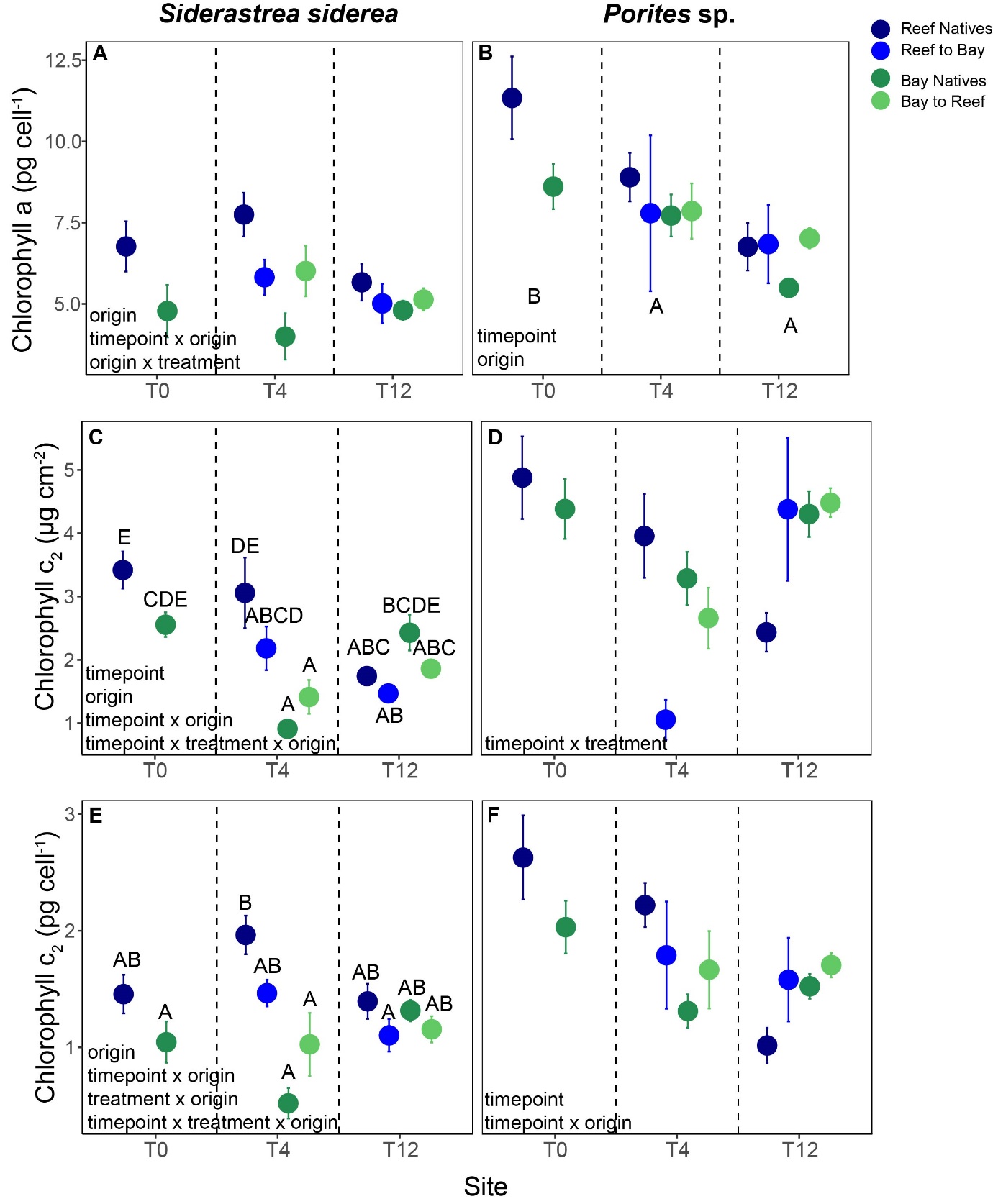

**Figure S5**

Chlorophyll a concentrations per cell (A-B), area-normalized chlorophyll c_2_ concentrations (C-D), and chlorophyll c_2_ concentrations per cell (E-F) for the four transplant groups of *Siderastrea siderea* and branching *Porites* sp. after 0, 4 and 12 months of transplantation (T0, T4, T12). Error bars are standard error. Shown is mean ± SE. Significant main effects and interactive terms are indicated in bottom left of each panel. Letters represent *post-hoc* results of pairwise comparisons for three-way interactions only (see Table S9 for all *post-hoc* results).
